# Single Cell Multiomics Across Nine Mammals Reveals Cell Type Specific Regulatory Conservation in the Brain

**DOI:** 10.1101/2025.08.06.668931

**Authors:** Ashlyn G. Anderson, Brianne B. Rogers, Erin A. Barinaga, Jacob M. Loupe, Elisa WaMaina, S. Quinn Johnston, Henry L. Limbo, Elizabeth A. Gardner, Anna J. Moyer, Amanda L. Gross, Douglas R. Martin, Summer B. Thyme, Lindsay F. Rizzardi, Richard M. Myers, Gregory M. Cooper, J. Nicholas Cochran

**Affiliations:** HudsonAlpha Institute for Biotechnology, Huntsville, AL, USA; University of Alabama at Birmingham, Birmingham, AL, USA; Department of Biochemistry and Molecular Biotechnology, The University of Massachusetts Chan Medical School, Worcester, MA; College of Veterinary Medicine, Auburn University, Auburn, Alabama, USA; Department of Biochemistry and Molecular Biology, The University of Alabama in Birmingham, Birmingham, AL, USA

## Abstract

Understanding the gene regulatory mechanisms underlying brain function is crucial for advancing knowledge of the genetic basis of neurologic diseases. Cis-regulatory elements (CREs) play a pivotal role in gene regulation, and their evolutionary conservation can offer valuable insights. Importantly, the function and evolution of CREs are affected not only by primary sequence, but also by the *cis*- and *trans*-regulatory context. However, comparative functional analyses across species have been limited, leaving how these regulatory landscapes evolve in the brain largely unresolved. Here, we generated single-nucleus multiomic (snRNA- and snATAC-seq) data from cortex tissue across nine mammalian species and identified candidate CREs (cCREs) in a cell type–specific manner. We developed a multidimensional framework of conservation to assess sites of shared function that integrates sequence, chromatin accessibility, and enhancer–gene associations. Using massively parallel reporter assays (MPRA) in human neural progenitor cells and neurons, we measured activity of cCREs including both conserved and human-specific regions. CRISPR interference validated conserved enhancer function, including at neurodevelopmentally important genes like *FAM181B*. Motif enrichment identified transcription factors distinguishing conserved versus recent cCREs. Linkage disequilibrium score regression indicated that both conserved and human-specific cCREs were enriched for neuropsychiatric GWAS risk, while neurodegenerative risk was confined to conserved elements. Our findings define functional dimensions of enhancer conservation and demonstrate how regulatory evolution shapes human brain biology and disease susceptibility.

## Introduction

The human brain has undergone substantial evolutionary change, including an expanded neocortex, increased cortical neuron number, and prolonged developmental timing^1, 2^. These changes are thought to arise in large part from genetic modification in regions that regulate gene expression, highlighting the importance of understanding how cis-regulatory elements (CREs), including enhancers and promoters, have evolved in humans^3–5^. Comparative studies have shown that while some CREs are conserved across mammals, others have diverged rapidly in specific lineages, contributing to species-specific traits^6–9^.

One prominent example of regulatory evolution involves human accelerated regions (HARs), which are highly conserved elements that have undergone rapid sequence divergence in humans. These elements often act as neurodevelopmental enhancers and contribute to human-specific brain features^10–13^. Large-scale efforts such as the Zoonomia project have expanded this view by systematically identifying conserved and recent CREs across mammals, revealing thousands of candidate CREs (cCREs) with lineage-specific sequence^14, 15^. However, these studies have focused predominantly on primary sequence conservations in CREs, which obscures cell type-specific regulatory programs and may overlook functional conservation that is not reflected at the sequence level^16^. This limitation is especially relevant in the brain, where diverse neuronal and glial populations have distinct regulatory programs. Lineage-specific changes in CREs often underlie differences in cell type-specific transcriptional programs across species, facilitating evolutionary fine-tuning of gene expression^17^. Furthermore, CREs can retain activity despite sequence divergence, and conversely, sequence conservation does not guarantee regulatory function in a given context^6, 16, 18, 19^. These complexities are particularly important for understanding human brain disorders, as genome wide association studies (GWAS) have shown that risk variants for neuropsychiatric and neurodegenerative disease primarily occur in non-coding regions, particularly in CREs active in specific brain cell types^20–24^. Indeed, many disease-associated variants likely impact disease risk by altering transcription factor (TF) binding in CREs and consequently altering gene expression^25, 26^. Notably, non-coding variants may occur in deeply conserved or recently evolved CREs, with distinct implications for neurodevelopment or lineage-specific traits.

To better understand the evolution of CRE functions in the brain, we generated a single nucleus multiomics atlas of chromatin accessibility and gene expression in nine mammalian species and zebrafish. By integrating snATAC-seq and snRNA-seq data, we identified cell-type specific cCREs across species, and classified their conservation based on sequence alignment, chromatin accessibility, and predicted target genes. We functionally assessed the activity of conserved and human-specific cCREs using massively parallel reporter assays (MPRAs) in human neuronal progenitor cells (NPCs) and neurons, and validated deeply conserved CRE-gene relationships through CRISPRi experiments in neurons. We further investigated the regulatory logic of these elements by evaluating TF binding site contributions using chromBPNet *de novo* motif discovery and ChIP-seq data from human brain tissue. Finally, we evaluated the disease relevance of these elements through partitioned heritability analyses and experimental validation of candidate disease-risk variants. By mapping functional conservation of CREs, we establish a framework for interpreting non-coding genetic variation. Together, this work provides a multidimensional view of CRE evolution in the brain and reveals how different modes of conservation relate to regulatory function, speciation, and disease susceptibility.

## Results

### Single nucleus multiomic profiling across nine mammals

To investigate how gene regulatory programs evolve in the mammalian brain, we integrated single nucleus multiomic (10x Epi Multiome) data of chromatin accessibility (snATAC-seq) and gene expression (snRNA-seq) from cortex tissue of nine mammalian species (**Figure 1a**). We generated data from five species (rat, cat, rabbit, horse, and cow) and leveraged published multiomic datasets from Anderson et al.^21^ (human) and Zemke et al.^27^ (macaque, marmoset, and mouse; **Figure 1a**). In total, we considered data from a median of three individuals per species (range 1–15), capturing a median of 8,279 nuclei per sample (range 2,257–17,390), and a median of 22,272 nuclei per species (range 7,308–105,332; Extended Data Fig. 1a, Supplementary Table 1-2). To contextualize mammalian regulatory conservation against more diverged species, we also generated single nucleus multiomic data from zebrafish telencephalon (**Figure 1a**, Extended Data Fig. 2). While our focus remained on conservation across mammals, inclusion of zebrafish allowed identification of deeply conserved regulatory elements shared across vertebrates.

**Figure 1.**
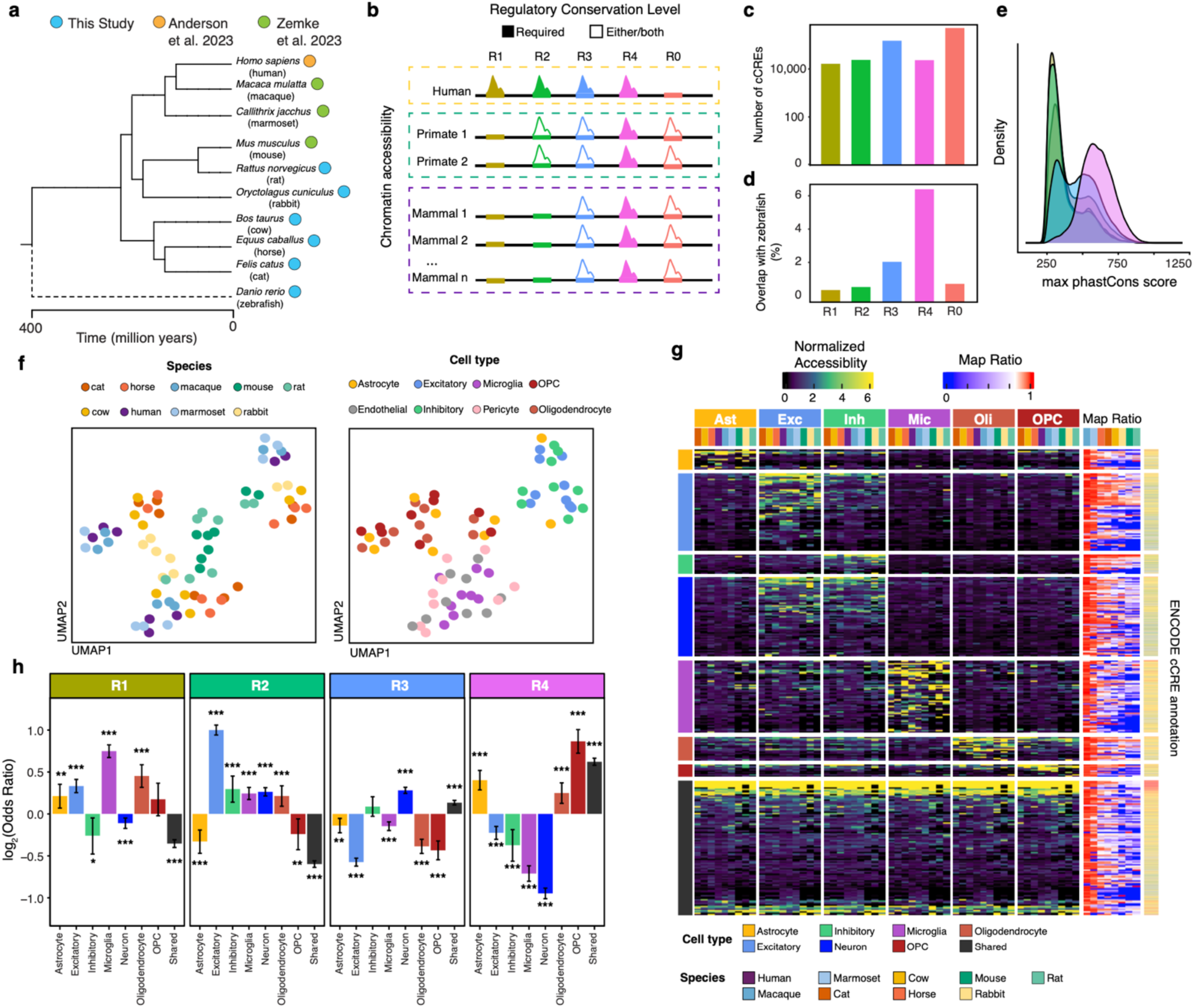
Regulatory conservation of brain cis-regulatory elements across mammals. **a)** Phylogenetic tree of the ten species included in this study. Circles indicate species with single nucleus multiomic data generated in this study or incorporated from prior studies. The dashed line indicates species used to evaluate deeply conserved vertebrate elements but not included in all analyses. **b)** Graphic illustrating the classification of CREs based on regulatory conservation: human-specific (R1), shared with primates (R2), shared with at least one non-primate mammal (R3), shared across all nine species (R4), and elements found only in non-human species (R0). **c)** Number of cCREs assigned to each regulatory conservation category. **d)** Proportion of cCREs in each category that overlap zebrafish cCREs, indicating deep vertebrate conservation. **e)** Sequence conservation of cCREs by regulatory conservation category, measured as the maximum phastCons score within each cCRE. **f)** UMAP of pseudobulk chromatin accessibility profiles across species and cell types, using peaks with orthologous sequences in all nine mammals. **g)** Left: heatmap showing pseudobulk accessibility profiles for randomized downsample of 15,000 cCRE within each cell type and species, grouped by the cell type-specificity of that cCRE as determined by the tau specificity index. Right: heatmap indicates the mapping ratio from hg38 to each species’ reference genome for these regions as a measure of sequence conservation. **H)** Enrichment of cell type-specific cCREs within each regulatory conservation category. Asterisks indicate statistical significance from Chi-squared test (*p < 0.05, **p < 0.01, ***p < 0.001).

We annotated cell types independently within each species by mapping snRNA-seq profiles to reference datasets using Azimuth^28–30^, enabling identification of eight broad cell type classes (astrocyte, endothelial cells, excitatory neurons, inhibitory neurons, microglia, oligodendrocytes, OPCs, and pericytes). We then integrated snRNA-seq across species on orthologous genes shared across all nine mammalian species and performed batch correction with Harmony^31^, confirming that cells clustered primarily by cell type rather than species and enabling confident identification of all major neuronal and glial populations (Extended Data Fig. 1b-j, Supplementary Table 2).

### Regulatory conservation of cCREs

To compare regulatory element function across species, we first identified cCREs by calling snATAC-seq peaks independently for each cell type within each species. We then lifted over snATAC-seq peaks from each species’ most recent genome assembly to hg38 coordinates, enabling direct comparison across species. We classified regulatory conservation for each cCRE based on the presence of an overlapping snATAC-seq peak in each species, resulting in five regulatory conservation levels: human-specific (R1), shared with primates (R2), shared with at least one non-primate mammal (R3), shared across all nine species (R4), and peaks found only in non-human species (R0; **Figure 1b**).

Across all species and cell types, we identified a total of 725,776 peaks, with an average of 217,501 peaks per species (range: 140,993–276,543; Supplementary Table 3, Supplementary Fig 3a,b). Of those, 213,367 peaks were detected in human. Categorization revealed that the majority of human peaks were conserved with respect to at least one non-primate mammal (R3 = 150,078); however, a small set of peaks were specific to humans (R1 = 16,427; **Figure 1c**, Extended Data Fig. 3a,b). To further identify the most deeply conserved cCREs, we overlapped our mammalian peaks with snATAC-seq peaks from zebrafish (n = 43,218). Although only 1.5% of peaks intersected zebrafish accessible regions, these were highly enriched for overlap with R4 peaks indicating that they are likely important to vertebrate biology (Chi-Squared test, OR = 5.32, p-value < 2.2x10^-16^; **Figure 1d**).

In addition, comparison to Zoonomia^14^ measurements of sequence conservation across 240 mammals at human cCREs defined by the ENCODE Project revealed that cCREs with high regulatory conservation in our data (R4) strongly overlapped the most conserved Zoonomia category (group-1; Chi-squared test, p < 2.2x10^-16^), whereas recently evolved elements (R1/R2) showed overlap across all Zoonomia conservation levels (Extended Data Fig. 3f). While cCREs with higher degrees of regulatory conservation exhibited increased sequence conservation by phastCons scores^32^ (Kruskal-Wallis test, p < 2.2x10^-16^), many cCREs shared between species do not exhibit elevated sequence conservation, emphasizing the importance of integrating functional conservation across species (**Figure 1e**, Extended Data Fig. 3c-e). Furthermore, recently evolved cCREs were enriched for transposable element-derived sequences, consistent with their established role in lineage-specific regulatory evolution^17, 27^, whereas conserved elements were depleted for transposable elements (Fisher test, OR= 2.64, adjusted p-value < 2.2x10^-16^, Extended Data Fig. 3g).

To examine cell type-specific patterns of regulatory conservation, we aggregated chromatin accessibility across all cells within each cell type to create pseudobulk accessibility profiles. Focusing on the subset of peaks that could be confidently mapped across all species, we performed dimensionality reduction with UMAP and correlation analyses. This revealed that accessibility profiles grouped predominantly by cell type, with closely related species grouping together within each cell type, suggesting that cell type-specific chromatin accessibility profiles are largely maintained within a cell type across mammals but exhibit a degree of phylogenetic specificity (**Figure 1f**).

### Conservation of cCRE cell type-specificity

Cell type-specific CREs are critical for directing cell type-specific gene expression. To measure cell type specificity of cCREs across species, we calculated a tau specificity index for each peak based on their pseudobulk chromatin accessibility profiles, where a score of zero indicates ubiquitous (across cell-types) accessibility and a score of one reflects complete cell type specificity^33^. Combining the tau score with fold change differences between cell types, we classified peaks as cell type-specific, pan-neuronal (shared between excitatory and inhibitory), or shared across multiple cell types (Supplementary Table 3). This classification was robust to cases where orthologous sequences existed only in a subset of species, enabling a comprehensive assessment of cell type-specificity independent of conservation status. When we examined accessibility patterns of cell type-specific peaks across species, we found they were largely consistent, with cell type-specific peaks maintaining selective accessibility profiles in the same cell types across mammals (**Figure 1g**).

To evaluate the accuracy of these classifications, we performed an orthogonal validation using publicly available sorted ChIP-seq data from more than 100 TFs and histone modifications profiled in the same human brain donors used for our single nucleus multiomics assays, including datasets from sorted NeuN+ (neuronal), Olig2+ (oligodendrocyte), and NeuN-/Olig2- (astrocyte and microglia-enriched) cell populations^34^. We observed that peaks classified as cell type-specific showed preferential overlap with ChIP-seq peaks from the corresponding sorted population (Extended Data Fig. 3h). Furthermore, only 6% of non-human peaks (R0) peaks overlapped any ChIP-seq peak from these datasets, supporting their lack of regulatory activity in human (Extended Data Fig. 3i).

We next asked whether certain cell type-specific cCREs were more likely to be deeply conserved or recently evolved. Microglia-specific cCREs were notably enriched among human-specific elements (R1; **Figure 1h**), suggesting more recent evolutionary innovation in this cell-type. In contrast, excitatory-specific cCREs showed the greatest enrichment among peaks shared across primates (R2), while peaks accessible across multiple cell types were most often found in the deeply conserved CREs (R4; **Figure 1h**). These findings indicate that while the regulatory programs underlying broad cell type identity are largely conserved, cell type-specific programs, particularly in microglia, have undergone recent evolutionary changes.

### Conservation of gene targets

Conservation of enhancer activity across evolution does not guarantee conservation of their target genes, as changes in three-dimensional chromatin architecture and large structural variants can reassign enhancers to new promoters, a process referred to as enhancer hijacking^12^. We mapped 346,339 cCRE-gene links across species, including 75,977 in human. Using our previous strategy, we classified target gene conservation into five categories: human-specific (T1), shared with primates (T2), shared with at least one non-primate mammal (T3), shared across all nine species (T4), and cCRE-gene links found only in non-human species (T0; **Figure 2a**). We found that a large portion of human cCRE-gene links were specific to humans (T1: 37.1%) with only a small fraction conserved across all nine species (T4: 0.59%; Supplementary Table 4). Even cCREs with high regulatory conservation frequently exhibited species-specific or partially conserved gene targets, highlighting that CRE–gene relationships are highly dynamic across mammals. Additionally, 37.8% of human links overlapped the cross species mammalian activity-by-contact (ABC) predictions from Zemke et al.^27^, which was based on single cell chromatin conformation, reflecting both the limited sensitivity of these methods and the complexity of regulatory interactions.

**Figure 2.**
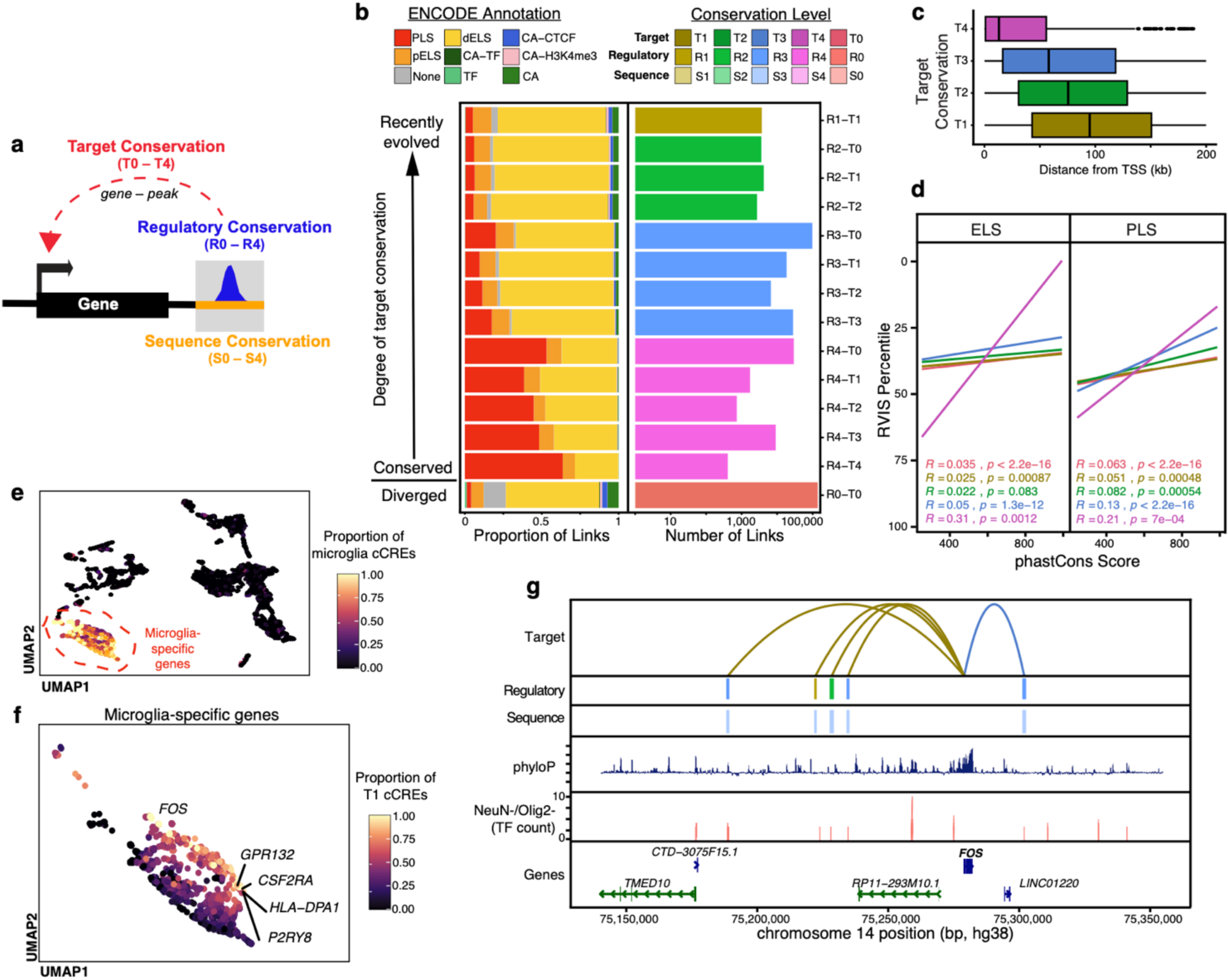
Conservation of predicted target genes across mammals. **a)** Diagram illustrating the three dimensions of conservation evaluated in this study: sequence conservation, regulatory conservation, and target gene conservation. **b)** Left: Proportion of cCRE-gene associations within each combined regulatory–target conservation category that overlap ENCODE cCRE annotations. Right: Number of cCRE-gene associations in each regulatory–target conservation category. **c)** Distance from cCREs to the linked-gene TSS by target gene conservation level. **d)** Pearson correlation between the sequence conservation of cCREs (phastCons score) and the constraint of their linked-genes (RVIS percentile), stratified by enhancer-like (ELS) versus promoter-like (PLS) cCREs. Positive correlations indicate that higher cCRE conservation is associated with genes under stronger selective constraint. **e)** UMAP embedding of genes based on the target conservation and cell type-specificity of all linked-cCREs for a gene, colored by the proportion of linked-cCREs for that gene that are microglia-specific. **f**) Zoomed-in UMAP of genes with the highest proportion of microglia-specific linked-cCREs, colored by the proportion of their cCRE associations that are human-specific (T1). **g**) Link plot of the *FOS* locus, with tracks showing (from top to bottom): target gene conservation, regulatory conservation, sequence conservation, phyloP scores, overlap with NeuN-/Olig2-brain ChIP-seq (number of TFs bound), and gene annotations.

When we overlaid ATAC-seq peaks with ENCODE-defined human CRE annotations^35^, links with the highest degree of regulatory and target conservation (R4–T4) more often consisted of promoter-like sequences (PLS, 63.6%; Chi-squared test, p < 2.2x10^-16^), while lineage-specific associations (R1–T1, R0–T0) were predominantly comprised of distal enhancer-like elements (dELS, 70.2% and 60.7%, respectively; **Figure 2b**). The distance to the transcription start site was significantly associated with target conservation (Anova, F = 1095, Df = 3, p = 2.2×10^-16^), with deeply conserved linked-peaks occurring markedly closer to their target genes than human-specific linked-peaks (**Figure 2c**). This association persisted even when restricting the analysis to distal enhancers, indicating that the association is not solely driven by promoter bias and is consistent with the idea that more proximal interactions are less likely to be disrupted by structural variants across evolution (Extended Data Fig. 4a).

We next examined how the sequence constraint of linked cCREs relates to intolerance of variation in their target genes. Using the Residual Variation Intolerance Score (RVIS)^36^, which quantifies whether genes harbor fewer or more common variants than expected given their expected mutational burden, we tested whether the maximum phastCons score of each linked-peak was associated with the RVIS percentile of the linked-gene. As expected, promoter sequence conservation was consistently correlated with a gene’s constraint across all target conservation levels (**Figure 2d**). At enhancers, this relationship was more nuanced. While the correlation (i.e., higher sequence conservation at cCREs correlating with higher degrees of intolerance of their target genes) remained significant across most target conservation categories, it was strongest for the most deeply conserved enhancer–gene associations (T4; **Figure 2d**). Using a linear interaction model, we confirmed that the slope of the relationship between cCRE phastCons score and gene RVIS percentile was significantly higher in T4 links compared to other target conservation levels (p < 0.0003 across comparisons; Extended Data Fig. 4b).

We investigated how the regulatory context of each gene varies by integrating both the cell type-specificity and evolutionary conservation of its linked cCREs. Using this information to create gene-level profiles and applying UMAP for dimensional reduction, we found that genes largely clustered by the cell type bias of their cCREs. Genes with a higher proportion of neuronal or shared cell type cCREs separated from those enriched for microglial or oligodendrocyte cCRE links (**Figure 2e**). This approach also revealed differences in conservation patterns, with some genes predominantly regulated by highly conserved cCREs and others showing a greater preference for recently evolved or human-specific elements (Extended Data Fig. 4c-f). Additionally, within the microglia-enriched genes, there were some genes with a higher proportion of links that were T1 (**Figure 2f**). Several genes displayed a high fraction of human-specific linked-peaks, including *FOS*, *CSF2RA*, *HLA-DPA1*, and *P2RY8*. These genes are implicated in microglial immune function, signaling, and synapse remodeling^37–39^. These findings underscore that even among genes expressed in the same cell type across species, the particular CREs regulating those genes can differ substantially between species.

To illustrate this, we examined the *FOS* locus in further detail. In humans, five microglia-specific cCREs were linked to *FOS*, of which four were human-specific cCRE–gene associations (T1). Surprisingly, three of these four were within cCREs exhibiting regulatory conservation across mammals (R2/R3), suggesting that while the cCREs themselves remain active across species, their association with *FOS* has been rewired in humans (**Figure 2g**). In non-human species, these cCREs were instead linked to, *MLH3*, and *JDP2*, with the *JDP2* association conserved across species (T3; Extended Data Fig. 4g). These linked cCREs also overlapped TF binding peaks identified in NeuN-/Olig2-nuclei, highlighting their activity in microglial-specific regulation.

### Transcription factor motif grammar across evolution

To investigate how TF motifs underlie patterns of cCRE conservation across cell types, we examined motif enrichments in snATAC-seq peaks grouped by regulatory conservation category (R1–R4) and cell type-specificity. We first calculated motif enrichments and scaled them across peak groups, then we performed hierarchical clustering to identify modules of co-enriched motifs (**Figure 3a**). This analysis revealed regulatory signatures associated with both cell type identity and conservation level. Motifs enriched in microglia-specific cCREs clustered together (module D) and were largely shared across conservation categories, including canonical microglial TFs such as SPI1 (PU.1) and RUNX1, consistent with their critical roles in microglia and immune gene regulation^40^ (**Figure 3a**, Supplementary Table 5). In contrast, we identified a subset of motifs predominantly enriched in R4 cCREs (module C), independent of cell type. These included broadly acting housekeeping TFs such as YY1, CTCF, and SP1, reflecting fundamental chromatin architectural programs with functions that are conserved across both cell-types and species.

**Figure 3.**
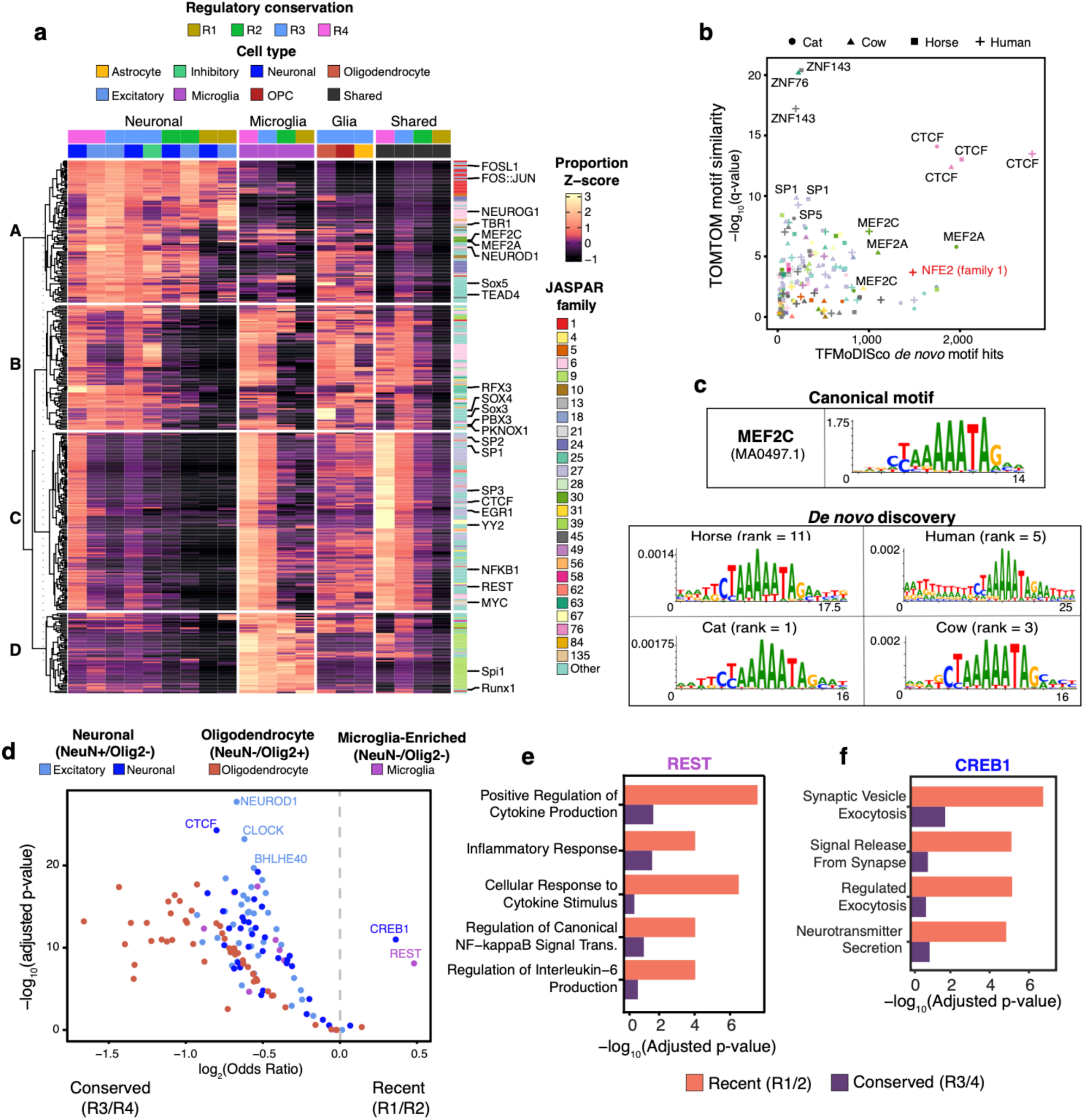
TF motif enrichment and binding reveal divergent regulatory contributions across conservation levels. **a)** Heatmap showing enriched JASPAR 2022 motifs across cCRE sets stratified by regulatory conservation and cell type specificity. Heatmap is colored by the row-scaled proportion of cCREs containing each motif. Rows split by k-means clusters (k=4) to identify modules of co-enriched motifs, with the right annotation indicating the JASPAR motif family. Columns are split by cell type specificity level. **b**) *De novo* motif discovery on NeuN+ ATAC-seq using chromBPNet to regress out enzyme bias. Scatter plot shows the number of occurrences of each identified motif (TFMoDISco) versus the TOMTOM similarity -log10(q-value) to known motifs, colored by the JASPAR motif family. **c**) Comparison of the *de novo* MEF2C motif recovered from NeuN+ ATAC-seq across species to the canonical JASPAR MEF2C motifs. **d**) Enrichment (Chi-squared test) of sorted brain TF ChIP-seq peaks in recent (R1/R2) versus conserved (R3/R4) cCREs, colored by the cell type specificity of the cCRE. cCREs were only intersected with ChIP-seq peaks from the corresponding sorted population. **e**) GO biological process term enrichment (hypergeometric test) for genes associated with recent versus conserved REST binding sites in microglia, using all genes linked to microglia-specific cCREs as the background. **f)** GO biological process term enrichment (hypergeometric test) for genes associated with recent versus conserved CREB1 binding sites in neurons, using all genes linked to excitatory- or neuronal-specific cCREs as the background.

The most notable divergence was within neuron-specific cCREs. We observed two distinct neuronal regulatory modules: one associated with deeply conserved neuronal cCREs (module B), including factors like PKNOX1, SOX3, and RFX3, which are involved in neuronal development and subtype specification, and a second, which had enrichment in all conservation categories, including human-specific cCREs (module A; **Figure 3a**). This cluster included activity-regulated and neurodevelopmental TFs such as FOS, NEUROG1, TBR1, MEF2C, and NEUROD1. This highlights the possibility that subtle shifts in neuronal enhancer grammar have contributed to human-specific changes in neuronal gene regulation.

To improve the ATAC-seq signal in neurons and generate data optimized for chromBPNet analysis, we generated NeuN+ sorted ATAC-seq data from the same tissues and used chromBPNet to perform *de novo* motif discovery on human, cat, horse, and cow cortex (Extended Data Fig. 5a-h). ChromBPNet is a base pair-resolution deep learning model that learns regulatory DNA sequence grammar while correcting for assay-specific enzyme biases, allowing for unbiased identification of TF motifs^41^. Across species, we recovered a similar profile of TF motifs. We further aligned these motifs with JASPAR 2022 motif families to account for similarity in motifs within TF families and identified 35 distinct families (Supplementary Table 6). Most motif families were observed in multiple species, supporting the existence of conserved neuronal regulatory grammar (**Figure 3b**). For example, MEF2-family motifs were consistently among the most detected *de novo* discoveries in all four species, reflecting their well-established role in activity-dependent gene regulation in neurons^42, 43^. Composite weight matrices (CWM) for the *de novo* motif further showed their similarity to the canonical human MEF2C motif across species (**Figure 3c**). Although the majority of motif families were shared, we identified five families that were only observed in one species. We observed that only ATAC-seq peaks in human NeuN+ cells were found to enrich for motifs from JASPAR family 1, which contains a diverse group of bZIP transcription factors, including AP-1 family members FOS::JUN, BATF, BACH, MAF, and NFE2. This motif family was the third most observed motif in the human data (**Figure 3b**, Supplementary Table 6).

To complement our motif analyses, we used publicly available cell type-sorted brain ChIP-seq data from Loupe et al.^34^. This provided a direct measurement of TF occupancy in human tissues, which is important as TFs are not always bound at motif instances in a given cellular context and can occupy regions that lack their canonical motifs^44^. We intersected these datasets with our snATAC-seq peaks to assess TF binding preferences across regulatory conservation levels. This revealed that binding of most TFs was enriched at conserved regulatory elements (R3/R4). However, we observed two notable exceptions, with CREB1 showing preferential binding at recent (R1/R2) neuron-specific cCREs, and REST enriched at recent (R1/R2) microglia-specific cCREs (**Figure 3d**).

We examined the functional roles of these TFs in recent cCREs (R1/R2) by using the associations between cCREs and their target genes then conducting a GO enrichment analysis. Genes associated with recent REST-binding sites in microglia were significantly enriched for pathways related to inflammatory responses, cytokine production, and NF-kappaB signaling, suggesting evolutionary changes in neuroimmune regulatory networks. In contrast, genes linked to conserved REST-binding sites in microglia showed no such enrichment (**Figure 3e**). Similarly, genes associated with recent CREB1-binding sites in neurons were specifically enriched for processes related to synaptic vesicle exocytosis, neurotransmitter secretion, and chemical synaptic transmission, whereas these pathways were not significantly enriched among genes linked to conserved CREB1 sites (**Figure 3f**). Additional analyses showed that CREB1-bound microglial cCREs were enriched for similar pathways at comparable levels across recent and conserved sites, whereas CTCF- and NEUROD1-bound neuronal cCREs were enriched for neuronal development and ion transport only at conserved elements (Extended Data Fig. 5i-k, Supplementary Table 7).

### Neuronal cCRE activity

To functionally evaluate deeply conserved and recently evolved cCREs, we performed a massively parallel reporter assay (MPRA) on 750 cCREs. These cCREs were prioritized based on multiple genomic features. We selected peaks called in neurons that also overlapped ChIP-seq peaks from human brain^34^, overlapped ATAC-seq peaks from iPSC-derived neurons^45^, did not overlap promoter annotations^35^, and intersected either deeply conserved (group-1) or recently evolved (group-3) sequence cCRE annotations from Zoonomia^14^ (**Figure 4a**). Additionally, we included promoters from genes highly expressed in neurons as positive controls and scrambled sequences with matched GC-content as negative controls. The nominated regions spanned a broad range of regulatory conservation, with the majority mapping to conserved classes (R3: 40.7%; R4: 40.8%). However, we also profiled more recent cCREs (R1: 6.3%; R2: 12.3%), as well as some non-human cCREs (R0: 1.7%; Extended Data Fig. 6a-b).

**Figure 4.**
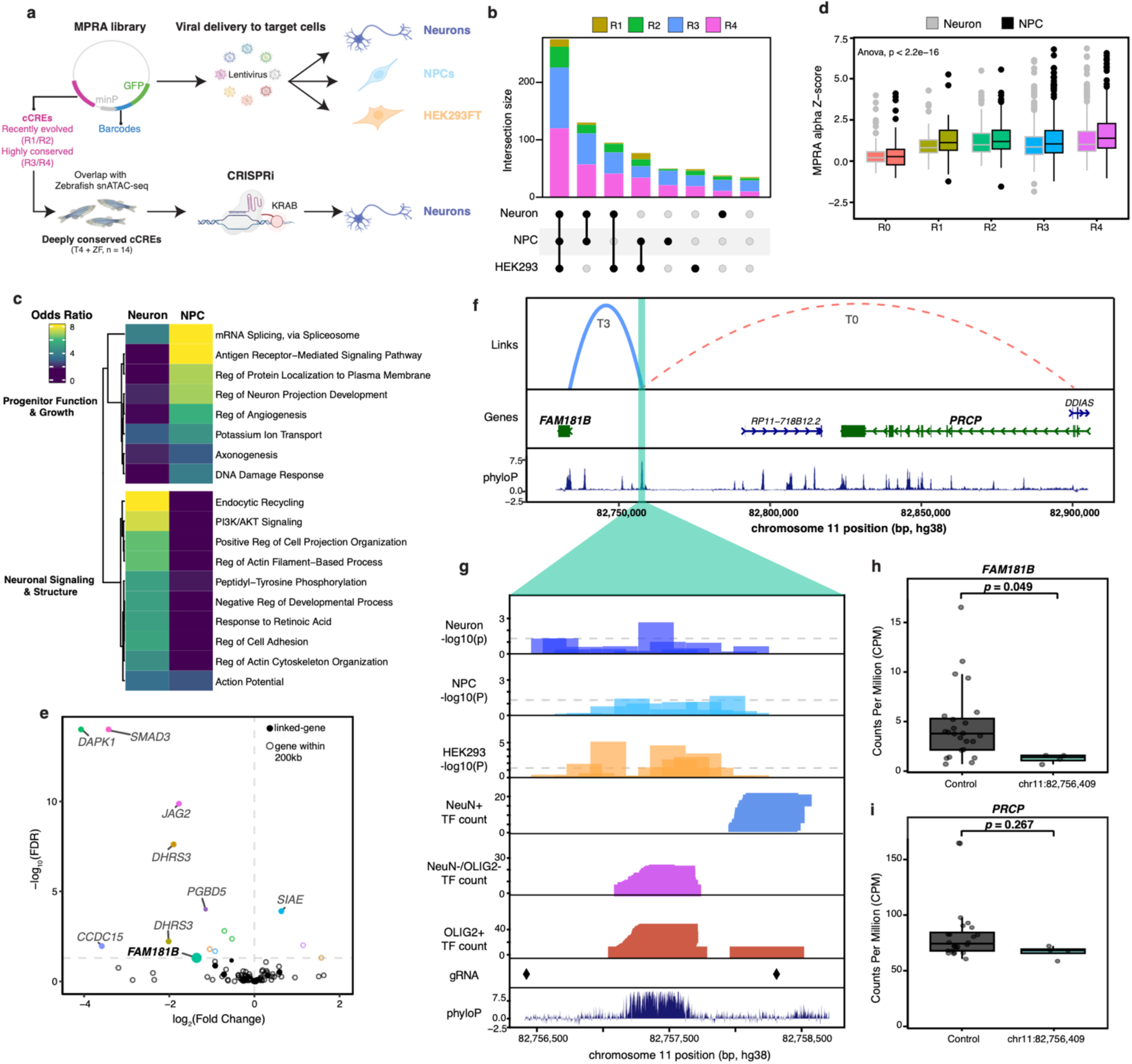
Functional validation of conserved and recently evolved cCREs using MPRA and CRISPRi. **a**) Schematic of the experimental design for functional validation, including MPRAs and CRISPRi. **b**) UpSet plot showing the number of regions with significant regulatory activity in the MPRA for neurons, NPCs, and HEK293FTs. **c**) Distribution of MPRAnalyze alpha scores by regulatory conservation category, separated by neuronal or NPC. **d**) GO term enrichment (hypergeometric test) for genes linked to regulatory elements with significant activity in NPCs (NPC, NPC & HEK293FTs) or in neurons (Neuron, Neuron & HEK293FTs). The background gene set includes all genes linked to regions tested in the MPRA. **e**) Volcano plot of CRISPRi DESeq2 results showing differential expression of genes within 200 kb of each targeted region. Dot colors indicate the specific targeted cCRE. **f**) Link plot of the *FAM181B* locus. Top plot: target conservation. Middle track: gene annotations. Bottom track: phyloP. **g**) Zoom in view of the targeted cCRE. Tracks from top to bottom show: -log10(p-value) from the MPRA in neurons, NPCs, and HEK293FTs; NeuN+, NeuN-/Olig2-, and Olig2+ TF binding counts; positions of gRNAs used for CRISPRi; and phyloP sequence conservation. **h**) Box plot showing expression of *FAM181B* in CRISPRi experiments comparing non-targeting controls to targeted cCRE perturbation. Value indicates the DEseq2 adjusted p-value. **i)** Box plot showing expression of *PRCP* for the same target cCRE. Value indicates the DEseq2 adjusted p-value.

We measured enhancer activity in HEK293FT cells, neuronal progenitor cells (NPCs), and iPSC-derived glutamatergic neurons. Most tested regions had significant regulatory activity in at least one cellular context, with almost 50% of regions exhibiting activity across all three cell types (**Figure 4b**, Supplementary Table 8). Furthermore, activity levels were significantly correlated between cell types (Extended Data Fig. 6c). However, activity was on average lower in neurons compared to other cell types (Extended Data Fig. 6d), potentially due to their post-mitotic state.

We also identified distinct sets of cCREs with activity predominantly in either NPCs or neurons (**Figure 4b**). Gene ontology analysis of genes linked to NPC-specific active cCREs revealed enrichment for mRNA splicing, axonogenesis, and neuron projection development, processes central to early neurogenesis^46^. Conversely, neuron-specific active cCREs were enriched for pathways such as endocytic recycling, PI3K/AKT signaling, and negative regulation of developmental processes, consistent with synaptic remodeling and terminal differentiation roles (**Figure 4c**)^47, 48^.

We found that recently evolved cCREs had activity levels similar to those that were deeply conserved (R1 vs R4; t-test, t = -1.27, Df = 907.94, p-value = 0.205; **Figure 4d**), suggesting that even recently evolved regulatory elements can acquire robust activity. In contrast, non-human cCREs (R0), which are accessible sites in other mammals but inactive in humans, had lower activity across human cell types (R0 vs R1-4, t-test, t=12.9, Df = 287.16, p-value < 2.2x10^-16^), consistent with functional differences of these sequences in the human lineage. When examining GO enrichment for genes linked to active cCREs, we found that recently evolved (R1/R2) cCREs were enriched for processes such as endocytic recycling and cell projection assembly, while conserved (R3/R4) cCREs were enriched for action potential, potassium ion transport, and neuron projection development (Extended Data Fig. 6e). However, the relatively small number of tested R1/R2 cCREs limits the strength of these conclusions.

### Validation of conserved enhancer-gene associations

To investigate the most deeply conserved neuronal cCREs, we prioritized R3/R4 cCREs that were also accessible in zebrafish and performed CRISPRi experiments in human iPSC-derived neurons. The inclusion of zebrafish data highlighted regions of deep vertebrate conservation. Focusing on non-promoter elements, we selected 14 deeply conserved neuronal enhancers (R3 = 8, R4 = 7; Extended Data Fig. 7a). For comparison, we also targeted the promoters of the genes linked to these enhancers, which were likewise conserved (R3 = 7, R4 = 10) though only 30% of these promoter regions overlapped accessible chromatin in zebrafish (Extended Data Fig. 7b). In total, we targeted 31 loci, comprising 14 enhancers and 17 promoters (**Figure 4a**, Supplementary Table 9).

Targeting these regions resulted in significant downregulation of at least one gene within 200 kb for 15 regions, including six enhancers and nine promoters, affecting the expression of 18 genes overall (**Figure 4e**, Extended Data Fig. 7c, Supplementary Table 10). Among these, 10 genes were directly linked to their targeted cCREs based on our cCRE–gene association maps defined using the single nucleus datasets. These validated regulatory relationships encompassed a range of target conservation levels, with most being shared across multiple species (T3 = 8, T2 = 1) but also including more lineage-specific interactions (T1 = 2, T0 = 3; Extended Data Fig. 7a). Among the most deeply conserved neuronal enhancers, we saw differential expression of *DHRS3, ROBO3, FAM181B, AHNAK2, SMAD3, PNKD,* and *CTDSP1* (Extended Data Fig. 7d-i).

Among the regions targeted in our CRISPRi experiments, one striking example was a deeply conserved neural enhancer linked to *FAM181B*. *FAM181B* is expressed in developing neural tissues and binds TEAD TFs, similar to co-regulators YAP and TAZ, and is a potential modulator of Hippo signaling in the nervous system, implicating it in neurodevelopmental growth control^49, 50^. The enhancer-gene association for *FAM181B* exhibited high target conservation across species (T3), and this cCRE was additionally linked to *PRCP* in three mammalian species, but not in humans (T0, **Figure 4f**). This cCRE demonstrated significant regulatory activity in our MPRA across all three tested cell types, supporting its broad regulatory activity (**Figure 4g**). The region also overlapped multiple TF binding sites from sorted human brain ChIP-seq data^34^. Interestingly, the positional footprint of TF binding differed by cell type, with TFs in NeuN+ nuclei binding approximately 500bp away from the glial-enriched populations (Olig2+, NeuN-/Olig2-), with binding for neuronal factors such as CREB1, NEUROD1, SATB2, TBR1, and MEF2C (Extended Data Fig. 7j). Additionally, the most conserved sequence aligned more closely with TF occupancy in glial cells, containing binding for TFs such as CTCF, YY1, SP1, and REST (Extended Data Fig. 7j). Perturbing this enhancer in neurons resulted in significant downregulation of *FAM181B* (log2FC = -1.36, adjusted p-value = 0.049; **Figure 4h**), whereas expression of T0-linked gene, *PRCP,* remained unchanged (log2FC = -0.26, adjusted p-value = 0.27; **Figure 4i**).

### Human lost CREs

We next sought to characterize cCREs that were deeply conserved across eight non-human mammals yet specifically lost in humans. This analysis identified 1,091 human-lost (HL) cCREs that were consistently accessible across all non-human species but exhibited no detectable chromatin accessibility or TF binding in any human brain cell type (**Figure 5a**, Supplementary Table 3). All HL cCREs could be lifted over to the human genome, indicating that their loss was not due to large sequence differences. In fact, sequence conservation was higher on average at HL cCREs than R3 conserved cCREs, though not as high as at the most deeply conserved R4 cCREs (t-test, HL vs. R3: p-value < 2.2e-16, HL vs. R4: p-value < 5.9x10^-9^, **Figure 5b**).

**Figure 5.**
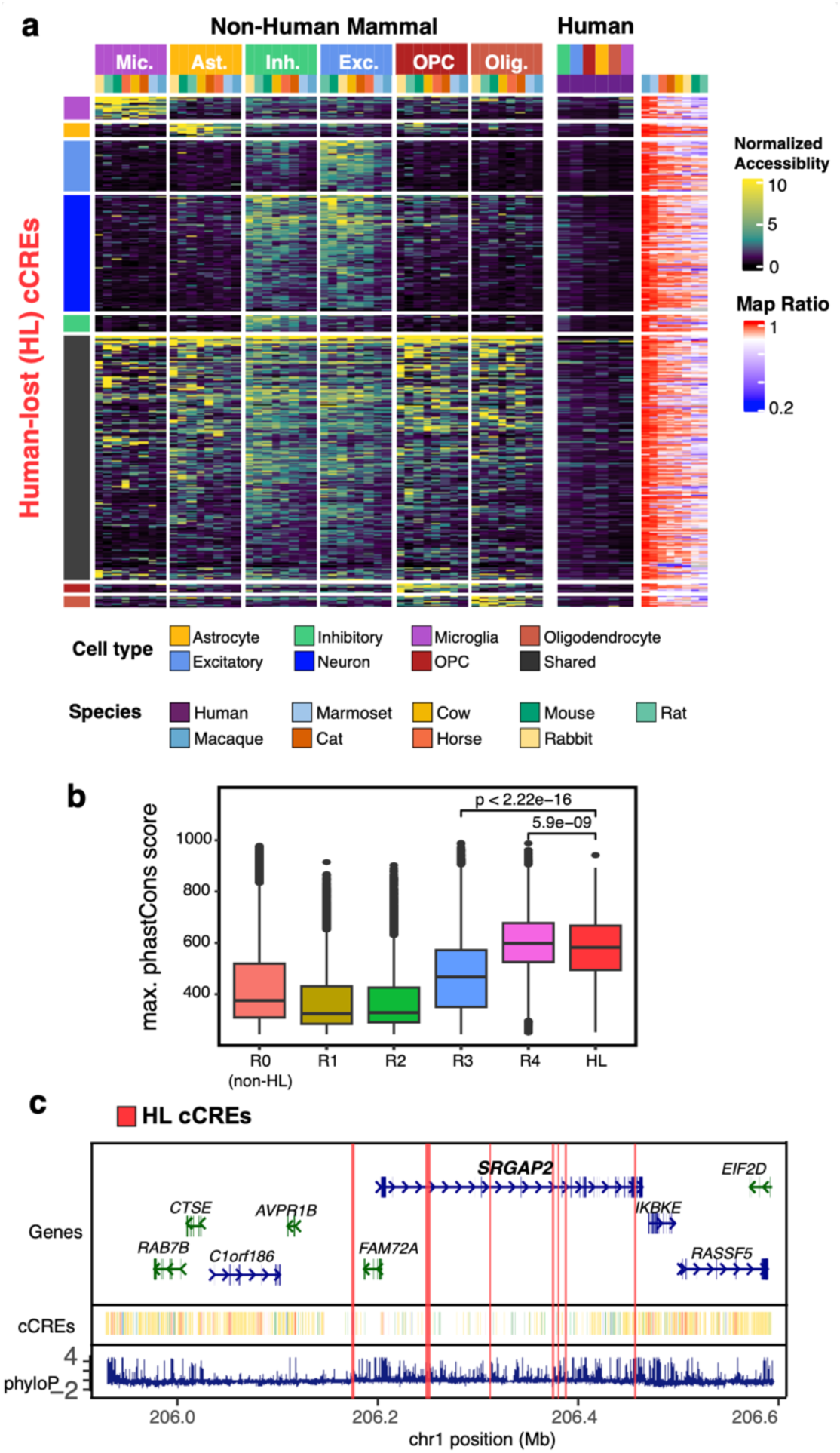
Characterization of human-lost cCREs. **a**) Heatmap of pseudobulk chromatin accessibility profiles across all cell type-species combinations for HL cCREs. Rows are split by the cell type specificity of the cCRE. The side heatmap indicates the mapping ratio from hg38 to each species’ reference genome for these regions. **b**) Maximum phastCons score in a cCRE by regulatory conservation level. Number indicates t-test p-value. **c**) Locus plot of *SRGAP2*. Top track shows gene annotations, middle track shows ENCODE cCRE annotations, and bottom track displays phyloP sequence conservation. Highlighted regions mark the locations of HL cCREs.

Consistent with this, HL cCREs were significantly enriched for overlap with zebrafish snATAC-seq peaks, indicating that many represent deeply conserved vertebrate CREs (Chi-Squared test, Odds Ratio = 1.58, p-value =0.025). Furthermore, only 3.1% of HL cCREs overlapped HARs^12^ and human-specific deletions in conserved regions (hCONDELs)^15^, highlighting that these regulatory losses are largely uncaptured by previous measures of human-specific sequence change in cCREs.

Cell type-specific analyses revealed that HL cCREs were significantly enriched among excitatory- and inhibitory-specific cCREs (Fisher test, Excitatory OR =2.09, adjusted p-value = 2.89x10^-101^; Inhibitory OR = 2.75, adjusted p-value =4.75x10^-7^), while shared cCREs were significantly depleted (Fisher test, OR = 0.65, adjusted p-value = 6.68x10^-11^), indicating that human regulatory losses preferentially impacted cell type-specific gene regulation (Extended Data Fig. 8a). Gene set enrichment analysis of the genes linked to HL cCREs showed enrichment across diverse biological processes. For example, microglia-linked HL cCREs were enriched for terms such as dendritic cell migration and chemotaxis, processes integral to microglial surveillance. Across cell types, linked genes also showed enrichment for categories including regulation of Wnt signaling and neural precursor cell proliferation (Extended Data Fig. 8b).

The gene linked to the most HL cCREs was *SRGAP2*, with nine HL cCREs. Of these, five were microglia-specific and four were shared across cell types (**Figure 5c**). *SRGAP2* has previously been implicated in the evolution of human-specific brain traits, but via gene duplications for *SRGAP2B* and *SRGAP2C*^51^. However, changes in the regulation of *SRGAP2* may have acted jointly with the gene duplication events to shape human-specific changes in synaptic density.

To assess whether similar patterns of regulatory loss occurred in other species, we performed a complementary analysis of mouse-lost cCREs. We identified 644 cCREs lost in mice but conserved across the other eight species. These mouse-lost cCREs were enriched in excitatory (Fisher test, Excitatory OR = 2.63, adjusted p-value =1.63x10^-11^) and oligodendrocyte-specific cCREs (Fisher test, OR = 2.42, adjusted p-value = 0.0013; Extended Data Fig. 8a). Mouse-lost cCREs were linked to genes such as *XKR4*, which is associated with gray matter volume^52^, and *PRTFDC1*, which has become an inactive pseudogene specifically in mice despite vertebrate conservation^52^. This parallel analysis highlights that lineage-specific regulatory loss is not unique to humans and may underlie species-specific brain features.

### Functional Conservation Stratifies Disease Heritability

We next performed linkage disequilibrium score regression (LDSC)^53^ to quantify how different combinations of functional conservation and cell type specificity contribute to the heritability of brain-related diseases. For this analysis, we focused on cCRE sets containing at least 1,000 elements to ensure robust heritability estimates relative to the background and used GWAS summary statistics from both brain-related and non-brain traits^54–71^ (Supplementary Table 11). As expected, LDSC coefficients were significant in cell types already associated with each disease, validating our cell type-specific annotations (**Figure 6a**, Supplementary Table 12). However, the degree of regulatory and target conservation for these enrichments varied across diseases. For example, Alzheimer’s disease (AD) heritability was predominantly enriched in microglia-specific cCREs, consistent with previous studies^72^, yet this enrichment was largely confined to conserved elements (R3 and T3; **Figure 6a**). In contrast, neuropsychiatric traits including bipolar disorder, neuroticism, and schizophrenia were significant across nearly all conservation categories. Notably, human-specific cCREs (R1, T1) showed significant contributions for these three disorders in neuronal cCREs, while conserved cCREs (R3, T3) were enriched both in neuronal and in shared cCREs spanning multiple cell types, suggesting that variation in both conserved and recently evolved cCREs contribute to polygenic risk for psychiatric disease (**Figure 6a**). Notably, autism spectrum disorder (ASD) heritability was most enriched at cCREs with regulatory conservation (R3) but human-specific gene targets (T1), suggesting that while the underlying cCRE activity is broadly conserved across mammals, the genes these cCREs regulate may have shifted in humans to influence ASD risk (**Figure 6a**).

**Figure 6.**
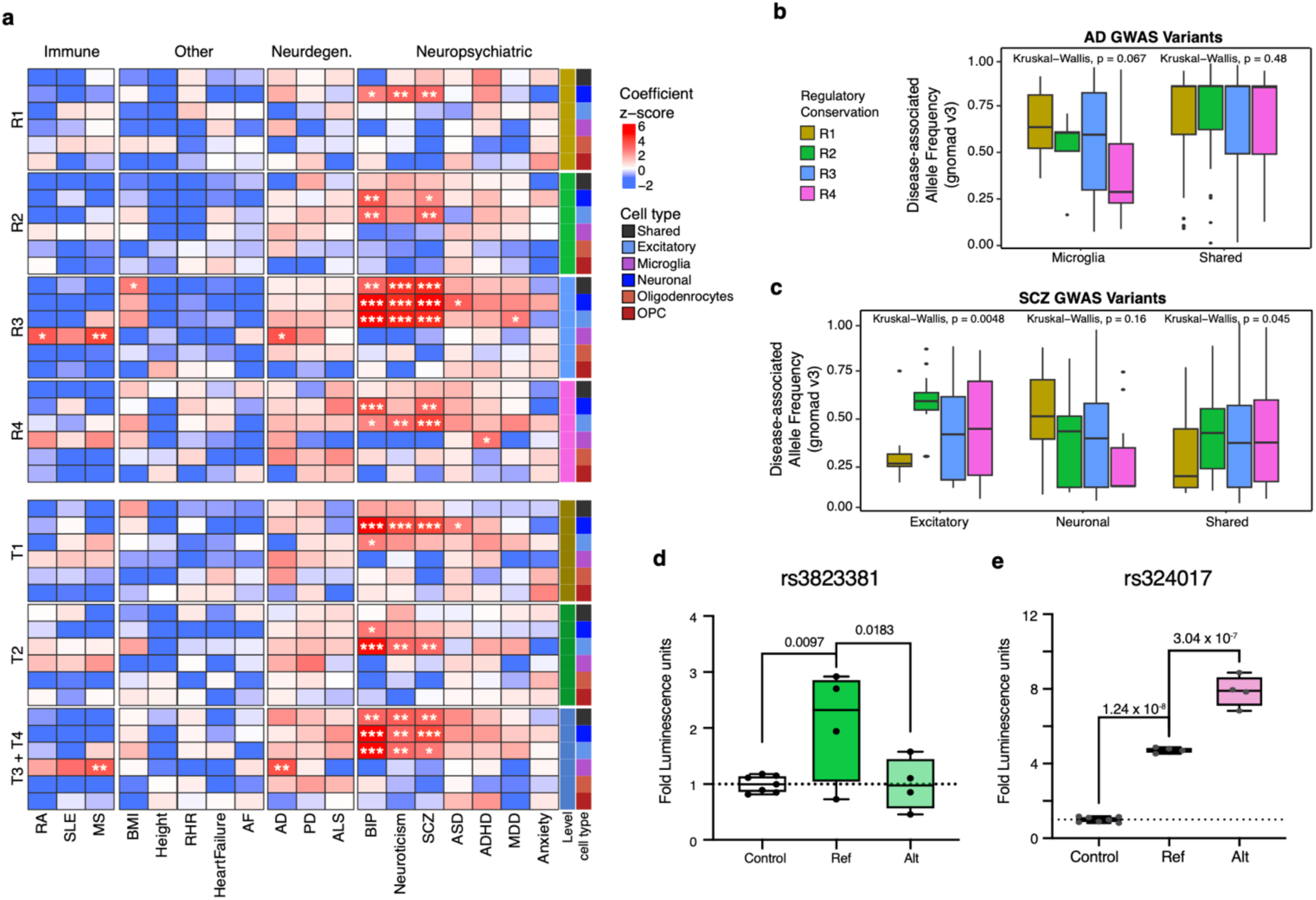
Functional conservation stratifies brain disease risk and heritability. **a)** Heatmap of LDSC heritability enrichment results for brain-related and non-related GWAS, shown for regulatory conservation levels (top) and target conservation levels (bottom). Columns are grouped by disease type. Asterisks indicate significance (* p<0.05, ** p<0.01, *** p<0.001). **b)** Distribution of gnomAD disease-associated allele frequencies for Alzheimer’s disease GWAS variants, stratified by the cell type specificity and regulatory conservation level of the cCREs they overlap. **c**) Distribution of gnomAD disease-associated allele frequencies for Schizophrenia GWAS variants, stratified by cell type specificity and regulatory conservation level of overlapping cCREs. **e**) Luciferase assay results showing the effect of selected schizophrenia-associated variant, rs3823381, on enhancer activity. P-value indicates significance of the coefficient in the linear model. **e**) Luciferase assay results showing the impact of selected schizophrenia-associated variant, rs324017, on enhancer activity. P-value indicates significance of the coefficient in the linear model.

The allele frequency (AF) of GWAS variants varied with regulatory conservation of cCREs in a cell type- and disease-specific manner. For Alzheimer’s disease (AD), microglia-specific cCREs showed a trend towards higher AF (gnomAD v3) of risk alleles in recently evolved elements (R1), while variants in highly conserved CREs (R4) exhibited lower AF, though this difference did not reach statistical significance (Kruskal-Wallis test, p = 0.067; **Figure 6b**, Extended Data Fig. 8c). In contrast, shared cCREs showed no significant variation in AF across conservation levels (p = 0.48). For schizophrenia, we observed a different pattern. Risk alleles in recently evolved excitatory cCREs (R1) had significantly lower AFs compared to more conserved elements (Kruskal-Wallis p = 0.0048). Neuronal cCREs overall did not show significant AF differences by conservation level (p = 0.16). However, shared cCREs displayed a modest but significant association, with lower risk allele frequencies in recently evolved elements (p = 0.045; **Figure 6c**).

Given our observation that schizophrenia, bipolar disorder, and neuroticism heritability is enriched across both recently evolved and conserved regulatory elements, we sought to directly test whether disease-associated variants in cCREs of different conservation levels could alter enhancer activity. We selected ten GWAS variants that were located within cCREs already shown to be active in HEK293FT cells in our MPRA data, including a range of conservation levels (R2 = 3, R3 = 6, and R4 = 1; Supplementary Table 13). Using luciferase reporter assays in HEK293s, we compared the regulatory activity of the reference versus the alternate allele. We found that the risk allele of rs3823381, a schizophrenia-associated variant located in a primate-specific cCRE (R2), modestly reduced enhancer activity (B = -1.07, p = 0.018, FDR=0.082), suggesting an allele-specific effect on gene regulation within a recently evolved CRE (**Figure 6d**). Additionally, we observed that the risk allele of rs324017, a schizophrenia-associated variant within a conserved cCRE (R3), significantly increased enhancer activity (B = 3.14, FDR = 2.74x10^-6^, **Figure 6e**, Extended Data Fig. 8d). This is consistent with previous reports showing that deletion of the rs324017 region in human iPSC-derived neurons led to downregulation of *LRP1* expression and altered electrophysiological properties, implicating this locus in neuronal excitability and schizophrenia-related phenotypes^73^. Together, these experiments demonstrate that risk variants in both recently evolved and conserved regulatory elements can impact enhancer function, supporting a model in which genetic perturbations across distinct evolutionary layers of gene regulation contribute to neuropsychiatric disease susceptibility.

## Discussion

In this study, we leveraged single nucleus multiomic profiling across nine mammals to develop a multidimensional view of CRE conservation in the brain. By integrating sequence, regulatory, and target gene conservation, we systematically characterized how both conserved and recently evolved cCREs shape cell type-specific gene regulatory programs. This work goes beyond traditional sequence conservation by revealing substantial conservation of regulatory activity even among cCREs with only modest sequence constraint and identifying functional divergence in target genes despite deeply conserved regulatory activity.

Functionally, we demonstrated via MPRA that both conserved and human-specific neuronal cCREs can drive robust enhancer activity in human neurons, NPCs, and HEK293FTs, emphasizing how recently evolved cCREs rapidly acquire functional roles. Complementing these findings, our CRISPRi experiments in neurons confirmed that deeply conserved enhancers regulate neurodevelopmental genes such as *FAM181B*, underscoring how fundamental regulatory relationships have been maintained across vertebrates.

Our analyses show that while most TFs preferentially bind conserved regulatory elements in humans, CREB1 and REST instead are enriched for binding at recently evolved cCREs in neurons and microglia, respectively. Notably, previous studies have shown that CREB1 binding sites were enriched for schizophrenia, bipolar disorder, and neuroticism heritability, while REST sites were enriched for Alzheimer’s disease risk^34^. These findings suggest that human-specific regulatory changes involving CREB1 and REST may underlie distinct vulnerabilities to neuropsychiatric and neurodegenerative diseases.

Our results highlight *FOS* regulation as an example of evolutionary changes in the human brain. We found that four out of five microglia-specific cCRE-*FOS* links are human-specific, pointing to substantial changes in how this gene is regulated in microglia. FOS is not only a marker of neuronal activity but is also expressed in glial cells, including astrocytes and microglia, where it is linked to proliferation and inflammatory pathways^75–77^. In microglia, FOS responds to lipopolysaccharides, paraquat, and excitotoxic conditions by driving cytokine production and inflammatory responses^75, 78–80^. Increased *FOS* expression is also seen in Alzheimer’s disease, where it is associated with amyloid accumulation, apoptosis, and cognitive decline^76^. Our findings suggest that human-specific changes in *FOS* regulation in microglia may have helped shape neuroimmune interactions in the human brain, which could also contribute to vulnerability to neuroinflammation and neurodegeneration. Future work that tests the impact of human-specific FOS function will be key to understanding how evolutionary changes in this regulatory network have influenced human brain function and disease risk.

Our study also revealed 1,091 cCREs that are active in all other tested mammalian species but not humans.These include nine such sites affecting *SRGAP2*, a gene that has previously been implicated in neurodevelopment and that has also experienced human-specific gene duplication events^51^. These HL sites had little overlap with previous annotations of sites with human-specific regulatory changes, indicating that our approach is unique relative to these efforts. Further, they are unlikely to be artifacts. For example, the fact that they were detected in all non-human mammals tested, and in many cases also zebrafish, and exhibited high degrees of mammalian sequence conservation suggests they are CREs that perform highly conserved functions. Conversely, their absence from 15 different human samples, coupled to the fact that they exhibited no TF binding in human brain tissue, makes it highly likely that these CREs are genuinely inactive in humans. Future analyses to determine the ancestral functions and gene targets of these CREs, the specific sequence changes in humans that have led to loss of function, and the biological consequences of these changes are likely to be of considerable interest. More generally, these findings are consistent with the “less is more” hypothesis that loss of function can lead to evolutionary novelty^81^, in this case contributing to human-specific alterations to the development and function of the brain.

Finally, our LDSC analyses provide insight into how regulatory conservation relates to brain disease risk. We show that Alzheimer’s disease heritability is predominantly enriched in conserved microglial cCREs, whereas neuropsychiatric traits such as schizophrenia, bipolar disorder, and neuroticism are influenced by both recently evolved and conserved neuronal cCREs. Furthermore, we found that heritability within recently evolved cCREs was predominantly confined to cell type-specific cCREs, whereas heritability within conserved cCREs spanned multiple cell types. Complementing these genome-wide patterns, our luciferase assays revealed that schizophrenia-associated variants in a primate-specific cCRE (rs3823381) and a deeply conserved *LRP1*-regulating CRE (rs324017) both altered enhancer activity. These results support a model where evolutionary changes in gene regulation, including both recent and conserved elements, have contributed to risk for neuropsychiatric disease.

This study has several limitations. First, cCRE–gene links were inferred based on correlations between chromatin accessibility and gene expression, and as such represent predictions rather than direct measurements of regulatory interactions. Confidence in these predictions decreases with increasing genomic distance, making distal associations particularly susceptible to false positives. Additionally, the power to detect cCRE–gene associations varies across species depending on sample size, potentially biasing comparisons of target conservation. While our analyses uncover broad patterns, functional validation of target gene associations was only performed for a subset of cCRE-gene links. Additional perturbation studies will be necessary to validate cCRE–gene associations. Second, the evolutionary distance between mammals and zebrafish presents a gap in our ability to resolve changes in regulatory conservation across vertebrates, which could be addressed by incorporating other species within this evolutionary gap in future work.

Overall, our work defines new principles of regulatory conservation in the brain and establishes a framework for interpreting non-coding genetic variation. Together, these findings provide a foundation for future studies to explore how the evolutionary changes in gene regulation contribute to human-specific cognitive traits and disease risk.

## Methods

### Brain tissues

Whole Sprague-Dawley rat and New Zealand White rabbit brains (1 male/ 1 female each) were obtained from BioIVT. For each sample, the hippocampus was first removed, then the brain was dissected along the longitudinal fissure to obtain the left and right cerebral hemispheres. To enrich for cortex tissue, the two hemispheres were further dissected by removing all tissue posterior to the corpus callosum. Horse and cow cortex samples were collected following euthanasia and necropsy from the Auburn University College of Veterinary Medicine necropsy service. Animals were kept at 4°C until organ collection, where cortex tissue was dissected and frozen on dry ice. Cat cortex tissues were obtained under Auburn University Institutional Animal Care and Use Committee (IACUC protocol 2018-3319) and comply with the Animal Welfare Act. Cats were sacrificed by intracardicac pentobarbital overdose followed by cold saline perfusion. The brain was divided into 6 mm coronal blocks and preserved via flash freezing in liquid nitrogen. Zebrafish experiments were approved by the UAB Institutional Animal Care and Use Committee (IACUC protocols 22155/21744). EK strain zebrafish were maintained on a 14 hour:10 hour light:dark cycle at 28°C. Twelve-month-old, female animals were anesthetized in 0.2% tricaine and rapidly euthanized by immersion in ice water before forebrain dissection. Samples were frozen on dry ice.

### Bulk RNA isolation and sequencing

For processing frozen brain tissue, all instruments and materials were chilled on dry ice and kept cold. At least 10 - 20 mg of frozen tissue was obtained from each sample using a metallic block and hammer to take a portion of the dissected cortex tissue. The tissue was then sealed inside of a Covaris Tissue TUBE (Covaris, cat. Nos. 520021 and 520023) and thoroughly pulverized using the chilled hammer and metallic block, with intermittent submerging into liquid nitrogen to maintain cold temperature and brittleness. Using a clean, pre-chilled scoopula, pulverized tissue was then transferred to a pre-chilled 2 mL DNA LoBind tube. RNA was isolated using the miRNeasy Mini Kit according to manufacturer instructions (QIAGEN, cat. no. 217084). RIN scores were calculated by a 2100 BioAnalzyer (Agilent) using the RNA 6000 Nano Kit following manufacturer instructions (Agilent cat. no. 50671511). At least 100 ng total RNA per sample was sent to Novogene for Eukaryotic mRNA library preparation with polyA enrichment. Samples were sequenced on a NovaSeq X Plus PE-150 flowcell, targeting 20 million reads per sample.

### Nuclei isolation from cortex tissue

The buffers required were nuclei extraction buffer (NEB: 0.32 M sucrose, 5 mM CaCl2, 3 mM Mg(Ac)2, 0.1 mM EDTA, 10 mM Tris-HCl, 0.1 mM PMSF, 0.1% Triton X-100, 1 mM dithiothreitol (DTT); Protector RNase Inhibitor (RI, 0.2U/µL) added before use according to manufacturer recommendation), sucrose cushion buffer (SCB; 1.6 M Sucrose, 3 mM Mg(Ac)2, 10 mM Tris-HCl, 1 mM DTT), interphase buffer (0.8 M Sucrose, 3 mM Mg(Ac)2, 10 mM Tris-HCl), blocking buffer (1xPBS, 1% BSA (Sigma, cat. No. 3117332001), 1 mM EDTA) and pellet buffer (add 200 µL 1 M CaCl2 to 10 mL SCB).

Methods for extracting and sorting nuclei from postmortem brain are similar to previously published methods^82^. Here, approximately 100 mg of tissue was placed into a chilled 1 mL Dounce homogenizer containing 500 µL of NEB on ice and allowed to partially thaw to ease douncing (2–3 minutes). Nuclei were extracted by douncing with a ‘tight’ pestle 30 – 40 times until the tissue was homogenized. The douncer was then washed with 500 µL NEB, and the entire 1 mL of homogenized tissue was passed through a 70 µm strainer into a fresh 2 mL DNA LoBind tube on ice. A sucrose gradient was prepared in two ultracentrifuge buckets (Beckman Coulter, cat. no. 344057) by layering 1 mL of Interphase Buffer on top of 1 mL SCB. The nuclear homogenate was carefully layered atop the sucrose gradient, balanced with NEB, then ultracentrifuged at 28,000 rpm for 30 minutes at 4°C in a SW28 swinging bucket rotor (Beckman Coulter). Upon completion, tubes were inspected for a visible pellet of nuclei at the bottom of the tube. Debris at the interface was removed first by using a P1000 pipette, then by continuing to remove the remaining sucrose gradient while being careful not to disturb the nuclei pellet. The pellets were carefully resuspended in 1 mL cold PBS and combined in a fresh 1.7 mL DNA LoBind tube on ice. A subset of nuclei were then labeled with DAPI for counting on a Countess II (ThermoFisher). The nuclei were then centrifuged at 500 x g for 5 minutes at 4°C to remove residual sucrose. Pelleted nuclei were then resuspended in 500 µL cold PBS.

### Single nucleus multiomics

Transposition, nuclei isolation, barcoding, and library preparation were performed according to the 10X Genomics Chromium Next GEM Single Cell Multiome protocol CG000338 Rev E with the following alterations. Each sample was loaded on one lane of the Chromium Next GEM Chip J. Nuclei were loaded according to manufacturer’s recommendations to target recovery of 10,000 nuclei per lane. For zebrafish, a sample was considered as a pool of 5 forebrain dissections due to tissue size and loaded on two lanes of the Chromium Next GEM Chip J. Libraries were sequenced by using Illumina NovaSeq X flowcells.

### Genome assemblies and annotation

*Homo sapiens* (human) assembly: hg38, GRCh38 annotation: hg38 Gencode v33; *Mus musculus* (mouse) assembly: mm10, GRCm38 annotation: mm10 Gencode vM22; *Macaca mulatta* (macaque) assembly: rheMac10, annotation: Ensembl release 110; *Callithrix jacchus* (marmoset) assembly: calJac4, annotation: Ensembl release 111; *Rattus rattus* (rat) assembly: mRatBN7.2 (rn7), annotation: Ensembl release 111; *Oryctolagus cuniculus* (rabbit) assembly: OynCun2, annotation: Ensembl release 111; *Bos taurus* (cow) assembly: bosTau9, annotation: GCF_002263795.2; *Equus caballus* (horse) assembly: equCab3, annotation: GCA_002863925.1; *Danio rerio* (zebrafish) assembly: danRer11, annotation: Ensembl release 109.

### Joint snRNA-seq and snATAC-seq workflow

Data was processed using cellranger-arc (v2.0.0). Low-quality cells were filtered on gene expression data (800 ≤ Number of genes ≤ 8000, and mitochondrial percent < 5) and chromatin accessibility data (ATAC fragments > 200, and ATAC fragments < 98% percentile). RNA counts were normalized with SCTransform^83^ with mitochondrial percent per cell regressed out. Principal component analysis (PCA) was performed on RNA, and UMAP was run on the first 30 principal components (PCs). The ATAC counts were normalized with term-frequency inverse-document-frequency (TFIDF). Dimension reduction was performed with singular value decomposition (SVD) of the normalized ATAC matrix. The ATAC UMAP was created using the 2^nd^ through the 50^th^ LSI components. Doublet density was computed using *computeDoubletDensity* from scDblFinder^84^ where doublet score is the ratio of densities of simulated doublets to the density in the data. Cells with a doublet score > 3.5 were removed. Normalization and dimension reduction were performed again on the filtered set with the same parameters.

For non-human mammals, predicted cell types were determined for each cell using Seurat SCT-normalized reference mapping. Gene expression data was mapped to SCT-normalized data and annotated with cell types using Azimuth^28^. For species annotated with a different species reference, cells with a predicted cell type score less than 0.75 were removed from the data. This threshold allowed for identification of broad cell types using the same cell type reference dataset while retaining species-specific cell populations.

For zebrafish, cells were clustered jointly using snRNA-seq and snATAC-seq (wknn) at a resolution of 0.3. Module scores for each cluster were calculated with Seurat’s *AddModuleScore* based on cluster markers from Pandey et al.^85^, and clusters were labeled according to the highest corresponding module score.

### Analysis of previously published snMultiome data

Raw fastqs were downloaded from GEO (GSE229169) for mouse, marmoset, and macaque from Zemke et al., 2023^27^. Each sample was first processed separately using *cellranger arc count*. Then samples from the same species were aggregated using *cellranger arc aggr*. All preprocessing and downstream QC was performed identically to datasets from the other species. Human data (GSE214979) was processed as previously described^21^.

### Gene-Peak association analysis

ATAC peaks were called independently for each cell type in each species using MACS2^86^ and Signac^87^ *CallPeaks.* The union of these peaks was used in downstream analyses. ATAC-seq peaks were filtered to only retain those with a -log10(Q-value) > 3. For each species, gene-peak associations were determined via the *cellranger-arc* (v2.0) *reanalyze* function using the cell type ATAC-seq peaks and only the nuclei that passed QC filtering. The maximum interaction distance was restricted to 200 kb. Gene-peak links with an absolute correlation score < 0.3 were removed.

### Cross species gene homology annotation

Gene orthologs were classified using Ensembl multiple species comparison tool (version 112), via biomaRt (v2.58.0)^88^. Ensembl gene IDs, whole genome alignment (WGA) scores, gene order conservation (GOC) scores, and sequence similarity were annotated for each species. All scoring metrics compare the ortholog to the annotated human gene. We included one-to-one, one-to-many, and many-to-many orthologs.

### Regulatory and target conservation classification

For each species, we identified orthologous human coordinates for snATAC-seq peaks using CrossMap^89^ and the appropriate UCSC chain file. Lifted-over regions were retained if they mapped with at least a mapping ratio of 0.25 and were excluded if the resulting human-mapped region exceeded 5 kb in width.

To define regulatory conservation across species, we used the hg38-lifted coordinates for snATAC-seq peaks from all species. Coordinates from each species were resized to 1 kb windows centered on the peak summit, and a consensus set of putative cCREs was generated by merging overlapping windows using the *reduce* function from the GenomicRanges package. Merged overlapping peaks larger than 2 kb were further split into non-overlapping 1 kb windows to maintain consistent resolution across the genome. Each consensus cCRE was then indexed and referenced back to the original peak calls in each species to determine presence or absence. The binary matrix of peak presence across species formed the basis for regulatory conservation classification. Peaks found only in human were designated R1. Peaks found in human and at least one non-human primate were classified as R2, and those found in human and at least one non-primate mammal were assigned R3. Peaks present in all nine species were classified as R4. Finally, peaks that were absent in human but had an orthologous human sequence in other species were classified as R0.

To assess target gene conservation, we first linked each cCRE to target genes within each species using cellranger arc to correlate chromatin accessibility and gene expression across cell types. To compare these links across species, we used Ensembl orthology annotations to identify the human ortholog for each gene in the non-human species. For each cCRE, we determined whether the orthologous cCRE was linked to the same orthologous gene. A cCRE was considered to have target conservation if it was linked to the same ortholog in another species. Target conservation levels were determined using the same strategy as regulatory conservation. Links found only in human were designated T1. Links found in human and at least one non-human primate were classified as T2. Those found in human and at least one non-primate mammal were assigned T3. Links present in all nine species were classified as T4. Links that were absent in human but had an orthologous human sequence and a human ortholog in other species were classified as T0.

### Human-lost cCRE classification

To identify human-lost cCREs, we classified consensus peaks as human-lost if they overlapped peaks called in all other mammalian species except human and showed no overlap with TF binding peaks from any cell type in human cortex. Mouse-lost cCREs were identified by lifting over peaks from each non-mouse species to the mm10 genome using CrossMap and the appropriate chain files. Overlapping peaks were defined using the same criteria applied in the human analysis. A cCRE was classified as mouse-lost if it overlapped peaks from all other mammalian species and did not overlap an H3K27ac peak in the mouse cortex (ENCFF008NDD).

### Sequence conservation classification

To assign levels of sequence conservation across species, we used CrossMap mapping ratios from hg38 to each species’ genome. Peaks that failed to lift over to any non-human species were classified as human-specific (S1). For all other peaks, we calculated the average and maximum mapping ratios across primate species and across non-primate mammals. Peaks were classified as ultra-conserved (S4) if they exhibited an average mapping ratio greater than 0.9 across both primates and non-primates. Peaks were classified as mammal-conserved (S3) if the maximum mapping ratio exceeded 0.25 in both primates and non-primates. Peaks were classified as primate-specific (S2) if the maximum mapping ratio exceeded 0.25 in primates but was below 0.25 in non-primate species. Peaks that did not meet these thresholds were labeled as variable. These mapping-based thresholds were chosen heuristically to reflect major patterns of evolutionary conservation across mammalian lineages.

### Cell type-specificity calculation

To assess the cell type-specificity of each CRE, we calculated the tau score based on normalized pseudobulk chromatin accessibility across species and cell types. For each species, we lifted over the full set of consensus peaks (1 kb windows in hg38 coordinates) to the species-specific genome assembly using CrossMap and the corresponding chain file. We retained only liftovers with a minimum mapping ratio of 0.25. Peaks that lifted over were retained regardless of whether they overlapped a called peak in that species.

For each cell type within each species, we pseudobulked accessibility by summing counts for each lifted-over region across all nuclei of that cell type. We then used the peak index to associate accessibility values across species back to their corresponding human coordinates, generating a matrix of pseudobulked accessibility values for each peak across all cell types and species.

The accessibility matrix was normalized using CPM(counts+1). Tau scores were then calculated for each cCRE across the merged, normalized accessibility profiles. The τ score quantifies specificity and ranges from 0 (ubiquitous accessibility) to 1 (cell type-specific accessibility), using the formula:

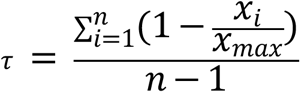

where *xi* is the normalized accessibility in cell type *i, xmax* is the maximum accessibility across all cell types, and *n* is the number of cell types.

Peaks were labeled as cell type-specific if τ > 0.7 and the log2 fold change between the most accessible cell type and the next highest was > 1.5. Peaks with τ > 0.5 where the two most accessible cell types were excitatory and inhibitory neurons were labeled as neuronal. All remaining peaks were classified as shared.

### Motif analysis

To identify TF motifs within snATAC-seq peaks, we called motifs using Siganc’s AddMotifs and JASPAR2022^90^ motifs for all peaks that lifted over to hg38. Motif enrichments were calculated using Signac’s FindMotifs for each cell type and regulatory conservation level, using 50,000 randomly selected, sequence-matched regions as the background. P-values were adjusted for multiple testing using the Benjamini-Hochberg method.

*De novo* motif discovery was performed independently for each species using chromBPNet^41^. NeuN+ ATAC-seq data from human^34^, cat, horse, and cow were used as input. For species with multiple samples, distinct samples were treated as replicates during preprocessing with the ENCODE pipeline. A matched background set was generated using “chrombpnet prep nonpeaks” for each species. Models were trained using fold 0 and the ENCSR868FGK_bias_fold_0.h5 bias model across all species.

Motif discovery was carried out using TFMoDISco on profile contribution scores from the bias-corrected chromBPNet models. Consensus contribution weight matrix (CWM) motifs were generated in both forward and reverse orientations. For each CWM, TOMTOM^91^ was used to identify the top three closest matches from a database of known MEME TF motifs along with common enzyme bias motifs. Significant matches were determined to be TF motif matches with a q-value < 0.0001, also supported by visual similarity to the CWMs.

### Partitioned heritability analysis

To evaluate whether feature links are enriched with common genetic variants that have been associated with brain-related GWAS, we performed stratified linkage disequilibrium (LD) score regression (sLDSC v1.0.1). sLDSC estimates the proportion of genome-wide SNP-based heritability that can be attributed to SNPs within a given genomic feature by a regression model that combines GWAS summary statistics with estimates of linkage disequilibrium from an ancestry-matched reference panel. Each cell type-conservation category was tested individually along with the full baseline model (baseline-LD model v2.2.) that included 97 categories capturing a broad set of genomic annotations. PMIDs for GWAS summary statistics are available in Supplemental Table 12. Additional files needed for the sLDSC analysis were downloaded from https://alkesgroup.broadinstitute.org/LDSCORE/ following instructions at https://github.com/bulik/ldsc/wiki.

### Gene set enrichment

The R package enrichR^92^ was used for gene set enrichment analyses when indicated. Gene sets were used as input to look for enrichment in GO Biological Process 2025, GO Molecular Function 2025, GO Cellular Component 2025, and KEGG 2025 databases. Terms with an adjusted p-value less than 0.05 were considered to be enriched. For gene set enrichment analyses using custom background sets, we implemented hypergeometric tests using custom scripts. Gene sets were obtained from the Enrichr website, specifically using the same GO collections. Enrichment was calculated relative to the appropriate background gene set for each analysis.

### Sorting Neuronal Nuclei

After centrifugation to remove sucrose gradient, nuclei were labeled by resuspending the pellet in 1 mL blocking buffer with NeuN-488 antibody (Millipore, cat. no. MAB377X) at 1:4,000 and 4,6-diamidino-2-phenylindole (DAPI) at 1:20,000 for at least 30 minutes at 4°C with rotation. Nuclei were then centrifuged at 500 x g for 5 minutes at 4°C. The supernatant was removed the pellet was resuspended in 500 μL cold 1% BSA in PBS with RNase Inhibitor. Nuclei were then filtered through a 35 μm filter, washing the filter with an additional 100 μL 1% BSA in PBS. Nuclei were then sorted using a Sony MA900 device with a 70-μm nozzle and pressure not exceeding a pressure setting of 7. Gates were set to capture those populations that were positive for 488 signal (NeuN^+^). Both the NeuN^+^ and NeuN^-^ populations were collected and held at 4°C. Purity of selected samples was typically >95% based on reanalysis of sorted samples.

### Sorted RNA-sequencing

Approximately 100,000 NeuN^+^ nuclei were sorted into 650 μL cold Trizol for RNA isolation. RNA isolation was performed using the Direct-zol RNA microprep kit (Zymo Research. cat. no. R2060) per manufacturer instructions. Sequencing libraries were prepared following the SMARTer Stranded Total RNA-seq Kit v2 – Pico Input (Takara Bio, cat. no. 634411). Using RIN scores from Bulk RNA preparations, a fragmentation time of 3 minutes was determined for each sample. Samples were sequenced on a single lane of a NovaSeq X Plus 10 B 300 cycle flowcell, targeting 150 million reads per sample.

### Sorted ATAC-sequencing

Approximately 100,000 NeuN^+^ nuclei were isolated from cortex tissues and used for ATAC-seq library preparation as per standard protocols^93–95^ with the following modifications. Briefly, 50 μL of cold 5x lysis buffer was added to sorted nuclei to reach a final concentration of 1x (1- mM Tris-HCl, pH 7.4, 10 mM NaCl, 3 mM MgCl2, 0.1% NP-40) and incubated for 20 min on ice followed by centrifugation for 10 minutes. The transposition reaction was incubated for 30 minutes at 37°C with agitation at 1000 rpm (Illumina Tagment DNA Enzyme and Buffer kit, cat. no. 20034198). After column clean up via the QIAGEN MinElute Reaction Cleanup kit (QIAGEN, cat. no. 28204) and PCR amplification of libraries, an additional clean up with AMPure XP beads (0.8X ratio) was performed with two 80% ethanol washes before quantification using a DNA High-Sensitivity chip on a 2100 BioAnalyzer (Agilent). Libraries were sequenced paired-end with the Illumina dual indexes on a NovaSeq X Plus 10 B 300 cycle flowcell targeting 100 million reads per sample. Samples were processed using the standard ENCODE ATAC-seq pipeline (https://github.com/ENCODE-DCC/atac-seq-pipeline v.1.7.0).

### Cell Lines

293FT cells were obtained from ThermoFisher Scientific (R70007) and maintained in DMEM (high glucose, L-glutamine, 100 mg/L sodium pyruvate) supplemented with 10% Fetal Bovine Serum (FBS), 1% Glutamax, 1% non-essential amino acids (NEAA), and 500 mg/mL Geneticin (G418, ThermoFisher Scientific). 293 cells were obtained from ATCC (CRL-1573) and maintained in EMEM (ATCC cat. no. 30-2003) supplemented with 10% FBS. KOLF2.1J Neural Progenitor Cells (NPCs) were generated from KOLF2.1J iPSCs, obtained from Jackson Laboratory, using the StemCell Technologies STEMdiff SMADi Neural Induction kit (StemCell Technologies cat. no. 08581). NPCs were maintained in NPC media: 2:1 DMEM (high glucose, L-Glutamine, 100 mg/L Sodium Pyruvate): Ham’s F12 Nutrient Mix and supplemented with 50X serum-free B-27 Supplement (ThermoFisher Scientific). hFG (40 μg/mL), hEGF (20 μg/mL), and heparin (5 μg/mL) were added daily for complete NPC media. KOLF2.1J-hNGN2 neurons were generated from KOLF2.1J-hNGN2 iPSCs obtained from Jackson Laboratory following a previously described protocol^45^.

### Plasmids

The psPAX2 (#12260), pMD2.G (#12259), FUGW-H1-GFP-neomycin (#37632), pLV hU6- sgRNA hUbC-dCas9-KRAB-T2a-Puro (#71236), and pLS-SceI (#137725) vectors were all obtained from Addgene. The pGL3.23 [*luc2*/minP] vector was obtained from Promega. Single-stranded 300 bp DNA oligos (n = 18,000) were generated by creating 270 bp sliding windows with 90 bp overlaps of candidate CREs with a max region of 2 kb and adding homology sequences for cloning. The sequences were generated by Twist Biosciences and cloned into the pLS-SceI vector, following the previously published LentiMPRA protocol^96^ with minor modifications as previously described^97^.

Luciferase elements were generated by selecting 467 bp of the nominated region and both the forward and reverse complement sequences were ordered as gBlocks from Integrated DNA Technologies (IDT). For cCREs with SNPs of interest, the variant allele was substituted for the reference allele. Elements were cloned in the the pGL4.23 vector digested with *EcoRV* by Gibson Assembly. Element insertion was confirmed by Sanger sequencing (MCLAB). Each element was individually prepped 3 times for a total of 6 individual plasmid preparations per nominated region.

CRISPRi sgRNA sequences were designed using https://benchling.com and ordered as premixed primer pools from IDT (Supp Table 9). The sgRNA sequences were then inserted into the pLV hU6-sgRNA hUbC-dCas9-KRAB-T2A-Puro plasmid by digesting with *Esp*3I and subsequent ligation, as previously described^98, 99^.

### MPRAs

MPRAs were performed in triplicate in HEK293FT cells, NPCs, and differentiated glutamatergic neurons following the Gordon et al. 2020^96^ protocol with minor modifications. HEK293FTs were plated at 70% confluency in two T225 flasks. The next day, HEK293FTs were transfected with 20 μg of oligo pools inserted into the pLS-SceI lentiviral vector per flask using Lipofectamine LTX with Plus Reagent. 48 hours after transfection, supernatant was harvested and filtered through a 0.45 μm bottle filter into a 100 mL bottle on ice. At this point, HEK293FTs used to produce lentivirus were harvested by treating with TrypLE Express for 5 minutes at room temperature, quenching with growth media, and cells pelleted by centrifugation at 200 x g for 5 minutes.

For NPCs, NPCs were plated at 50,000 cells/cm^2^ onto two matrigel coated 150 mm dishes per replicate. A replicate was considered an independent viral preparation and passage of NPCs. The next day, culture media was renewed with complete NPC media supplemented with protamine sulfate (10 mg/mL). To transduce NPCs, 11.25 mL of cold, filtered lentivirus was added to the NPC media. 24 hours post transduction, media was renewed with fresh complete NPC media. Cells were then incubated for four days before harvesting following the QIAGEN AllPrep DNA/RNA kit (cat. no. 80204).

For glutamatergic neurons, KOLF2.1J-hNGN2 iPSCs were plated at 25,000 cells/cm^2^ and differentiated following a previously published protocol^45^. On Day 14 of differentiation, media was renewed with Cortical Neuron Culture media supplemented with protamine sulfate (10 mg/mL). To transduce, 11.25 mL of cold, filtered lentivirus was added to the culture media. 24 hours post transduction, media was renewed with fresh Cortical Neuron Culture media. Cells were then incubated for four days before harvesting following the QIAGEN AllPrep DNA/RNA kit (cat. no. 80204).

Sequencing libraries were generated as previously described^96^ and sequenced at a 3:1 ratio of RNA:DNA on a NovaSeq X Plus 1.5 B 100 cycle flowcell targeting 150 M reads per replicate. Data was analyzed using MPRAflow as previously described^100^.

### Lentivirus Production

293FT cells were plated at 70,000 cells/cm^2^ in poly-L-ornithine coated 6-well culture plates. The next day, the media was renewed with OptiMEM Reduced Serum Media supplemented with 300 ng D-glucose. Cells were transfected with 1 μg pLV hU6-sgRNA hUbC-dCas9-KRAB-T2A-Puro plasmid with inserted gRNA sequence using Lipofectamine LTX with Plus Reagent, following manufacturer recommendations. 48 hours after transfection, supernatant was harvested and filtered through a 0.45 μm syringe filter into a 15 mL conical tube on ice.

### CRISPRi

NPCs were plated at 57,000 cells/cm^2^ in reduced growth factor Matrigel (Corning #354230)- coated 12-well culture plates. The next day, culture media was renewed with NPC media supplemented with protamine sulfate (10 mg/mL). To transduce NPCs, 500 μL of cold, filtered lentivirus was added to the NPC media and plates were centrifuged at 2,000 rpm for 30 minutes. 24 hours after transduction, cells were renewed with fresh NPC media with growth factors and 0.5 μg/mL puromycin for selection of successfully transduced cells. Cells were selected for 24 – 48 hours, until the control well (transduced with FUGW-H1-GFP-neomycin) had minimal remaining living cells. NPCs were then swapped to neuronal differentiation media following the Bardy et al. protocol^101^. Neurons were harvested for RNA isolation at day 14 of differentiation, following the Norgen Total RNA Purification kit (Norgen 37500) with the RNase-free DNase I kit (Norgen 25720). sgRNAs were designed upstream of the predicted target gene’s TSS in the promoter region as a positive control for each tested link region. sgRNAs targeting the *AAVS1* safe harbor locus were used as non-targeting controls. 100 ng total RNA per replicate was sent to Novogene for Eukaryotic mRNA library preparation with poly A enrichment. Samples were sequenced on a NovaSeq X Plus PE-150 flowcell, targeting 20 million reads per sample.

### RNA-seq analysis

Bulk RNA-seq data generated from tissues and CRISPRi experiments was analyzed as follows. Reads were trimmed using cutadapt for adapter removal (AGATCGGAAGAGC) and quality trimming (-m 30). Reads were then aligned to corresponding reference genome with STAR [-- outFilterType BySJout --outFilterMultimapNmax 20 --alignSJoverhangMin 8 -- alignSJDBoverhangMin 1 --outFilterMismatchNmax 999 --outFilterMismatchNoverReadLmax 0.04 --alignIntronMin 20 --alignIntronMax 1000000 --alignMatesGapMax 1000000 --outSAMtype BAM SortedByCoordinate --outSAMattributes NH HI NM MD]. Duplicate reads were filtered using picard MarkDuplicates. Counts were generated with htseq-count using the intersection-nonempty method and non-stranded counting. For sorted NeuN+ RNA-seq, only read-1 was used for data processing.

### CRISPRi Differential RNA-seq Analysis

The CRISPRi RNA-seq count matrix was normalized to counts per million (CPM). Each sample was then scored using Seurat’s AddModuleScore to determine the level of neuronal differentiation and astrocyte presence. Scores were calculated using a list of markers for differentiation (*CACNA1C, ENO2, MAP2*) and astrocytes (*AQP4, SLC1A3*), as previously described^45^. DEGs were determined for each target region versus non-targeting controls for all genes. Differential expression was calculated using DESeq2 with differentiation score and astrocyte score as covariates. Genes with an adjusted p-value< 0.05 were determined to be significant.

### Luciferase

293 cells were plated at 70,000 cells/cm^2^ in a 24-well format. The next day, cells were transfected with 1 μg of plasmid DNA using Lipofectamine LTX with Plus Reagent (ThermoFisher Scientific cat no. 15338-100) following the manufacturer’s recommendations. Per transfection 1 μg of luciferase element cloned into pGL4.23 [*luc2*/minP] were used, with pGL4.23 without inserted cCRE as a baseline control. Cell lysates were harvested by freezing at -80 °C 48 hours post-transfection. Luciferase assays were performed using the Bright-Glo Luciferase Reporter Assay System (Promega cat. no. E2610) following the manufacturer’s protocol. Cell lysis was performed on the 24-well plate and aliquoted across 4 wells of a white bottom 96-well plate for 4 technical replicates per biological replicate. Assays were completed in quadruplicate. Firefly luminescence was first normalized across the average plate luminescence then normalized to the average control luminescence. For each biological replicate, the median fold luminescence value was determined for the four technical replicates. At least two biological replicates were performed per region of interest. A linear model was used to assess enhancer activity of all reference alleles and to evaluate the effects of alternate alleles. Statistical significance was determined based on the t-statistics of the linear model coefficients, and Benjamini-Hochberg adjusted p-values are reported to account for multiple testing.

## Author Contributions Statement

Conceptualization: J.N.C Investigation: A.G.A, B.B.R, J.M.L., L.F.R, E.A.B, S.Q.J., H.L.L, E.W., E.G., Data Curation: A.G.A., B.B.R., Formal analysis: A.G.A, B.B.R., Provision of tissue: A.J.M., A.L.G., D.R.M., S.B.T., Supervision: R.M.M, G.M.C, J.N.C, Writing - Original Draft: A.G.A., Writing - Review and Editing: B.B.R., L.F.R, G.M.C, and J.N.C., Funding: J.N.C. and R.M.M.

## Acknowledgments

This work was funded by generous support from the HudsonAlpha Foundation, through the Clift Family Fund and the Leo Fund.

## Data availability

The raw and processed data generated are available through NCBI GEO under series accession number GSE303187.

## Code availability

The analysis scripts used to generate the images and statistical output in this paper are available on GitHub (https://github.com/aanderson54/snMulti_CrossSpeciesBrain).

## Extended Data

**Extended Data Fig. 1.**
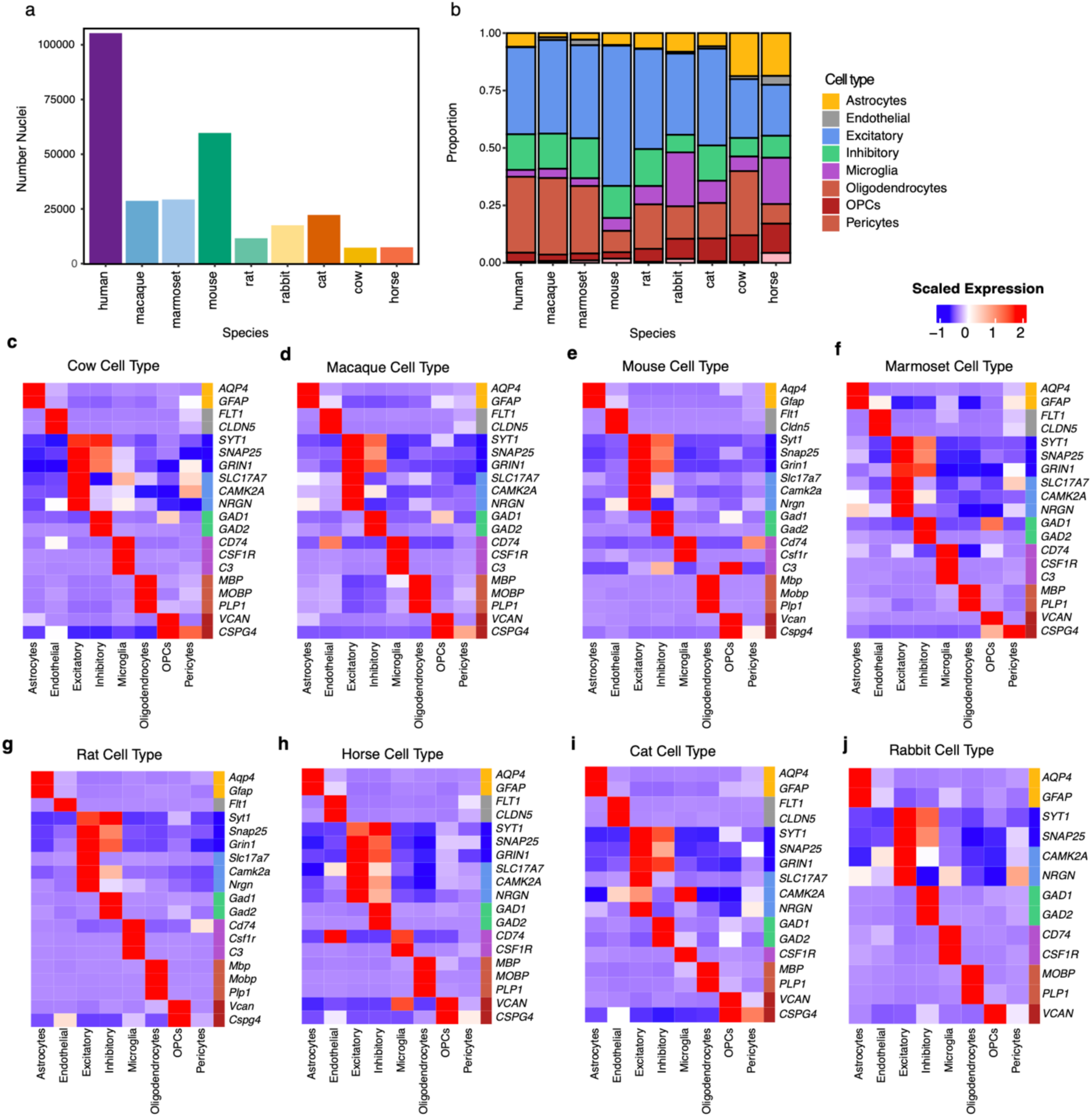
**a)** Number of nuclei profiled per species. **b**) Proportions of major cell types identified within each species. **c-j**) Heatmaps showing row-scaled expression of key cell type markers for cell types identified across all species. Only genes with annotated orthologs in Ensembl are shown for each species.

**Extended Data Fig. 2.**
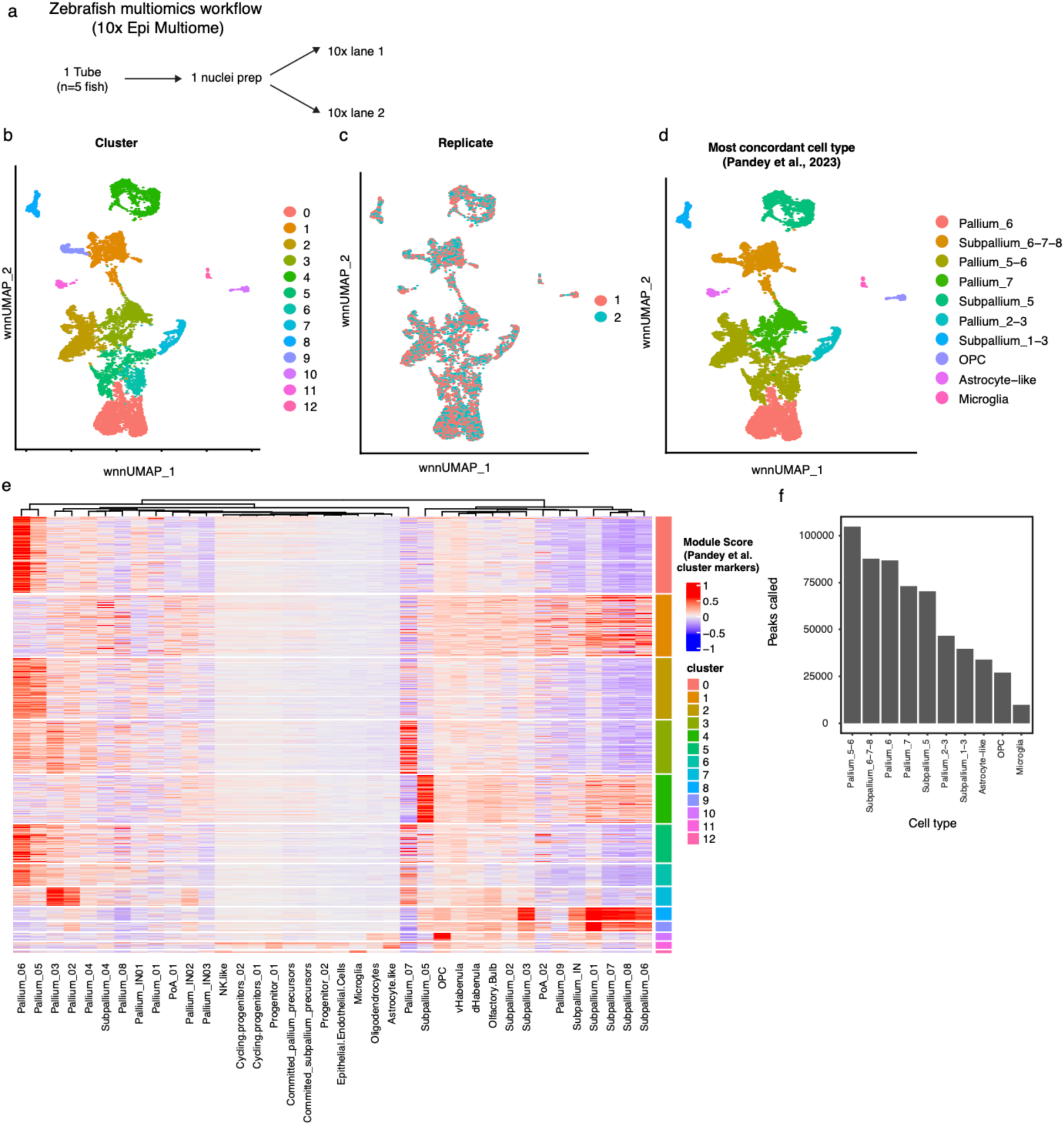
**a**) Schematic showing the workflow for zebrafish single nucleus multiome data generation. **b**) Weighted nearest neighbor (WNN) UMAP of zebrafish nuclei colored by cluster (resolution=0.3). **d**) WNN UMAP colored by 10x replicate to assess batch effects. **d)** WNN UMAP colored by most concordant cell type from Pandey et al. 2023 for each cluster. **e**) Heatmap of module scores for each cluster based on Pandey et al. cluster marker genes. **f**) Number of peaks called for each cluster in the zebrafish dataset.

**Extended Data Fig. 3.**
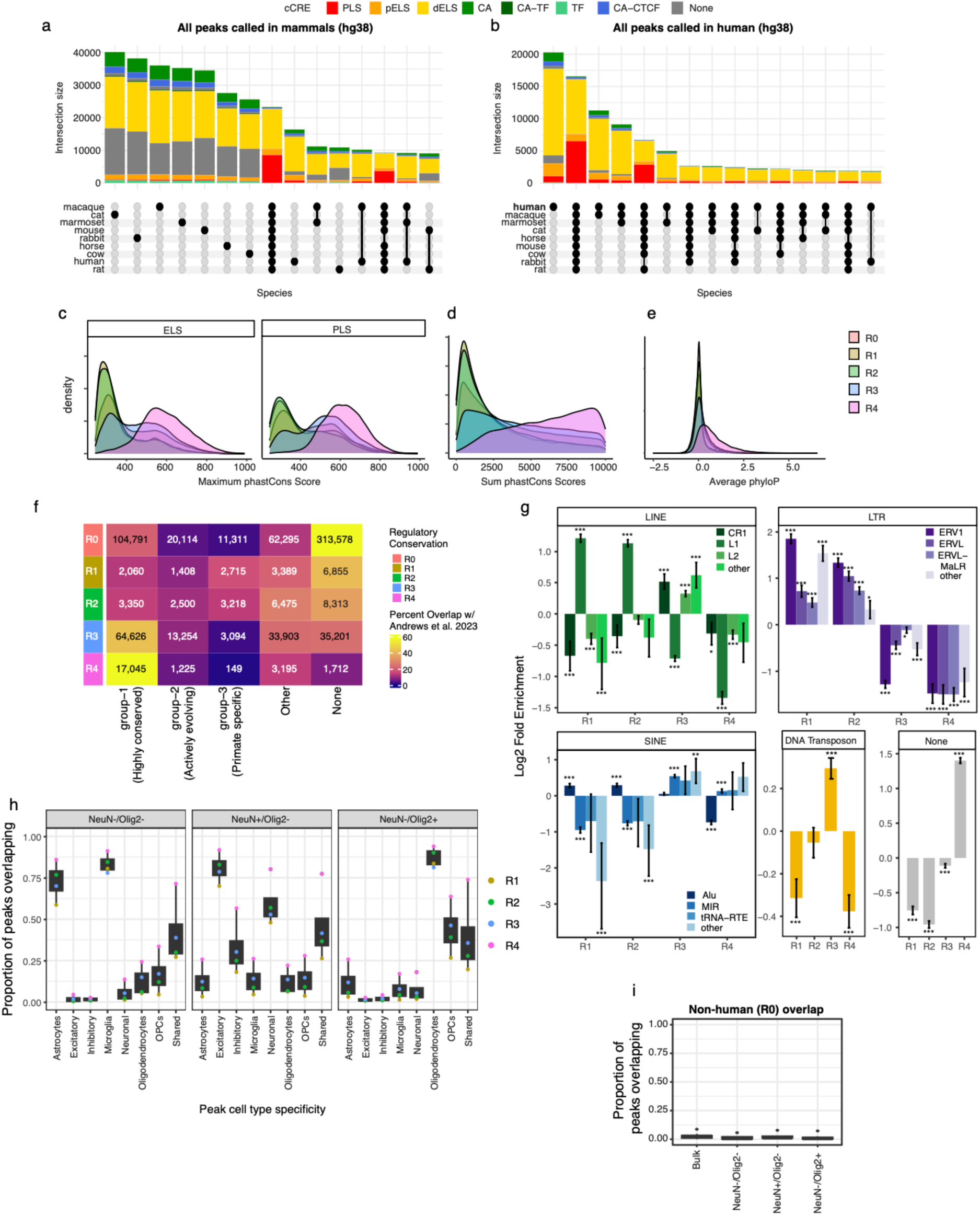
**a**) UpSet plot showing overlap of peaks between mammalian species for all peaks liftable to hg38. **b**) UpSet plot showing overlap of human peaks with peaks from other mammalian species. **c**) Density plot of maximum phastCons scores for cCREs by regulatory conservation level, split by promoter-like (PLS) and enhancer-like sequences (ELS). **d**) Density plot of summed phastCons scores for cCREs across regulatory conservation levels. **e**) Density plot of average phyloP scores for cCREs by regulatory conservation level. **f**) Heatmap showing the number of cCREs overlapping annotations from Andrews et al. by regulatory conservation level. **g**) Enrichment of transposable element overlap for cCREs by regulatory conservation level. Asterisks indicate Bonferroni adjusted p-values from Chi-squared test (*p < 0.05, **p < 0.01, ***p < 0.001). **h**) Box plots showing the proportion of cell type-specific cCREs overlapping sorted brain TF and histone ChIP-seq peaks from Loupe et al. **i**) Proportion of non-human-only (R0) cCREs that overlap the same ChIP-seq datasets.

**Extended Data Fig. 4.**
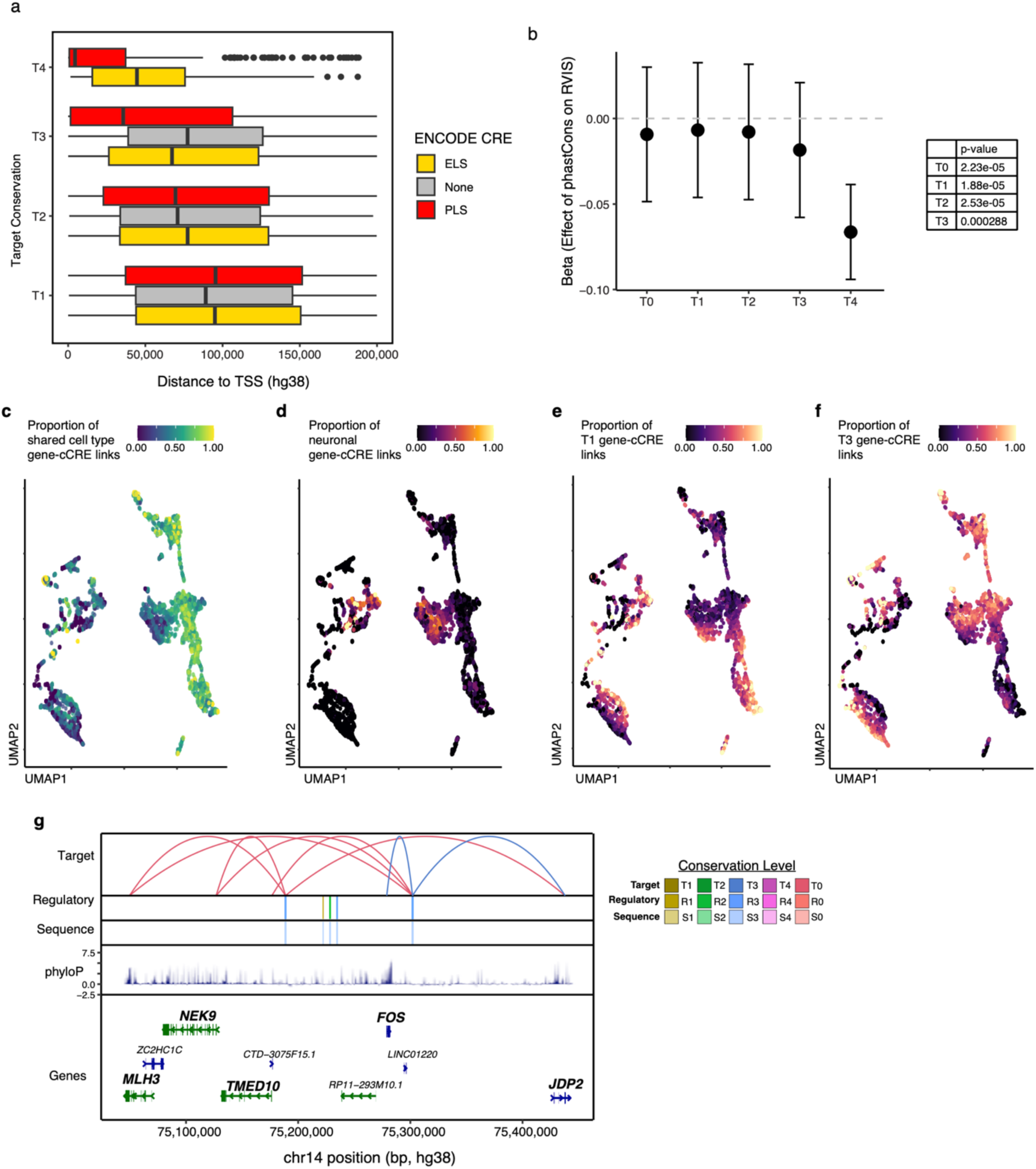
**a)** Distribution of distances from cCREs to the TSS of the linked-gene (hg38), stratified by target conservation category and overlap with ENCODE CRE annotations. PLS includes PLS and pELS annotations; ELS includes all other ENCODE regulatory categories; None indicates no overlap. **b**) Beta coefficients from a linear interaction model testing the association between linked cCRE phastCons scores and linked gene RVIS percentiles, by target conservation category. **c-f**) UMAPs corresponding to Figure 2E, colored by (**c**) shared cell type proportion, (**d)** neuronal proportion, (**e**) proportion of human-specific (T1) links, and (**f**) proportion of conserved (T3) links. **g)** Link Plot showing the *FOS* locus and non-human cCRE-gene links, with tracks showing (from top to bottom): target gene conservation, regulatory conservation, sequence conservation, phyloP scores, and gene annotations.

**Extended Data Fig. 5.**
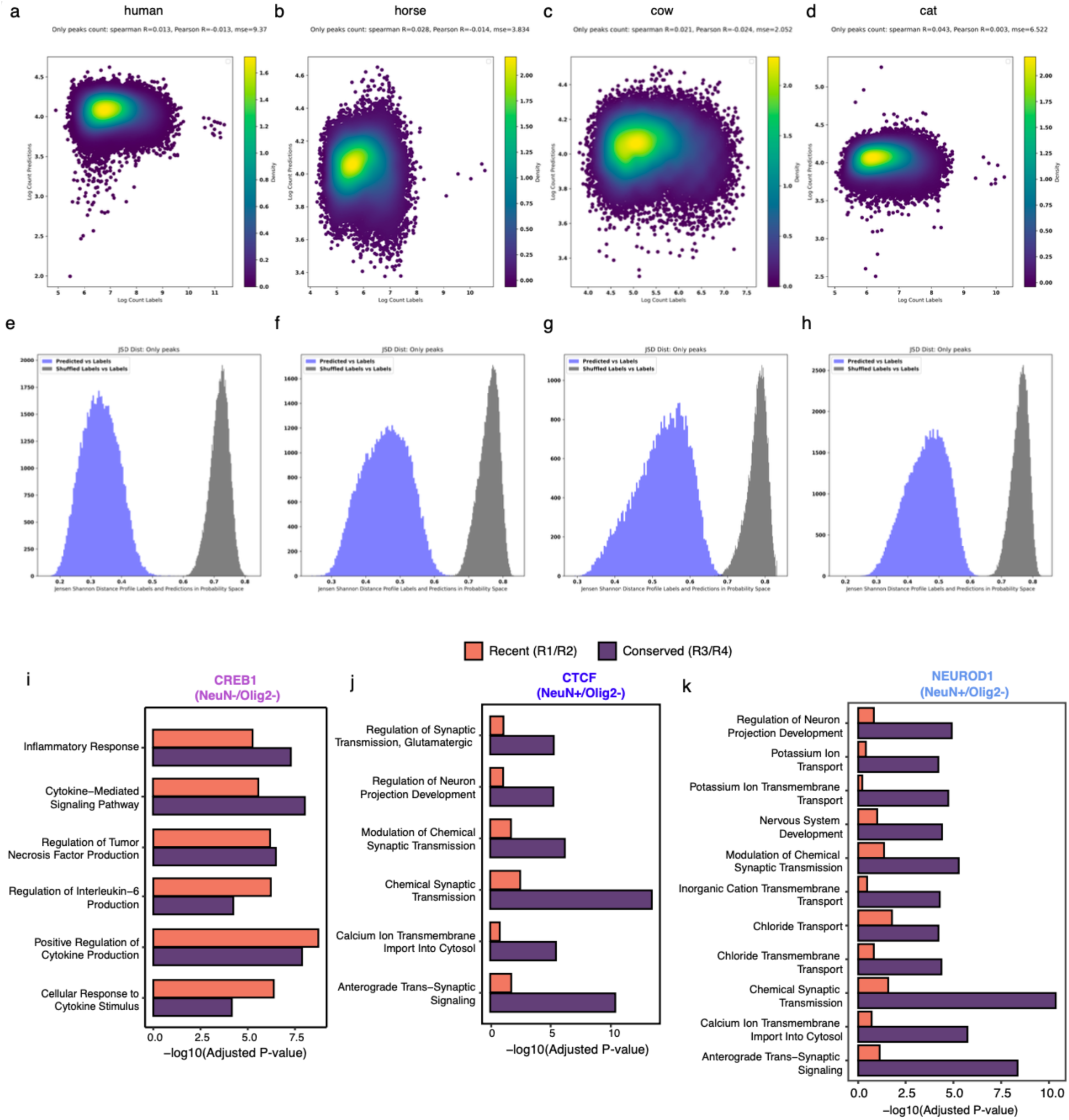
**a-d**) chromBPNet quality control plots for each species, showing the correlation between log-transformed observed counts and predicted counts for bias-only peaks. **e-h**) Additional chromBPNet QC, displaying the distribution of Jensen-Shannon distances between probability profiles for predicted and shuffled labels in each species. **i**) GO enrichment for genes linked to recent versus conserved CREB1 binding sites in microglia, with all genes linked to microglia-specific cCREs as the background. **j**) GO enrichment for genes linked to recent versus conserved CTCF binding sites at neuronal cCREs, relative to all genes linked to neuronal cCREs. **k**) GO enrichment for genes linked to recent versus conserved NEUROD1 binding sites in excitatory-specific cCREs, relative to all genes linked to excitatory-specific cCREs.

**Extended Data Fig. 6.**
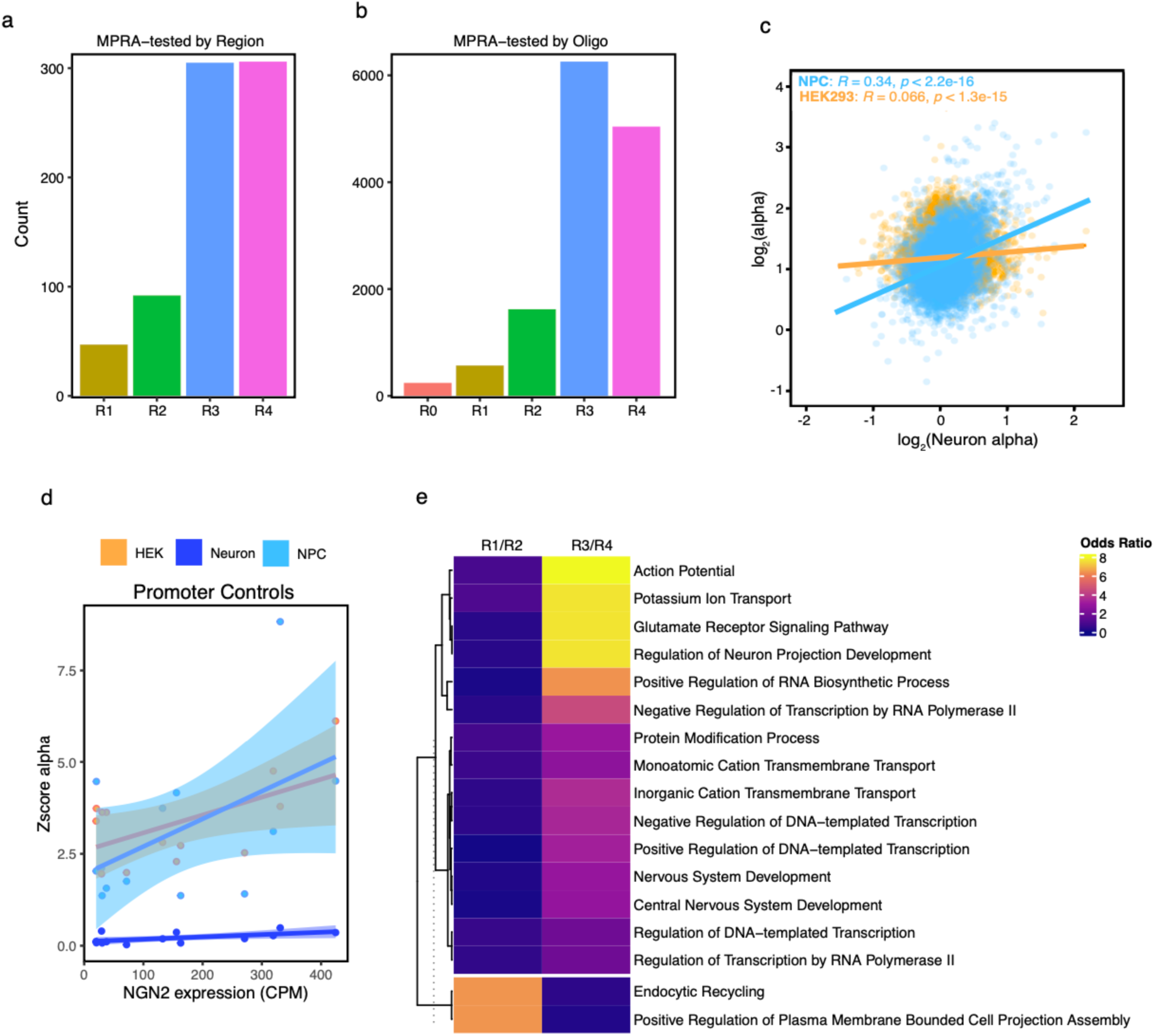
**a**) Number of tested regions in each regulatory conservation group, summarized at the region level. **b**) Number of tested oligonucleotides in each regulatory conservation group. **c**) Correlation of MPRAnalyze alpha scores between neurons and the other two cell types (NPCs and HEK293FTs). **d**) Scaled alpha scores for all promoter controls plotted against the expression of the corresponding gene in iPSC-derived neurons, colored by MPRA cell type. **e**) GO term enrichment for genes linked to active regions classified as recently evolved versus conserved, with all genes linked to tested regions serving as the background set.

**Extended Data Fig. 7.**
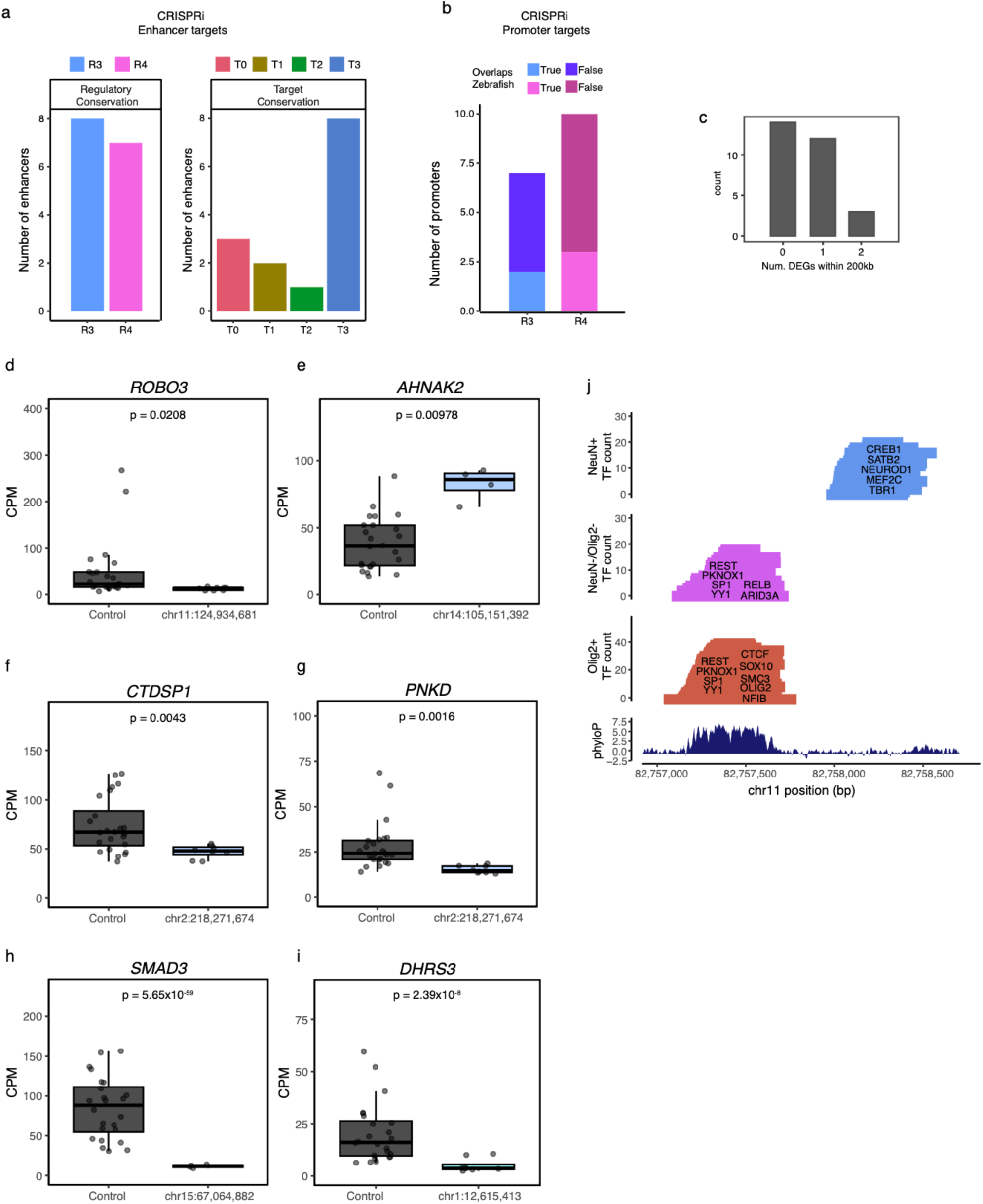
**a**) Number of enhancer regions tested in CRISPRi experiments by regulatory (left) and target (right) conservation level. **b**) Number of promoter regions tested in CRISPRi experiments by regulatory conservation level, colored by overlap with zebrafish chromatin accessibility. **c**) Number of differentially expressed genes (DEGs) within 200 kb of each targeted region. **d-i**) Box plots showing expression levels for non-targeting controls and targeted linked-CREs with DESeq2 adjusted p-values for (**d**) *ROBO3*, (**e**) *AHNAK2*, (**f**) *CTDSP1,*(**g**) *PNKD,* (**h**) *SMAD3*, and (**i**) *DHRS3*. **j**) Expanded locus plot corresponding to Figure 4g. Tracks from top to bottom show NeuN+ TF peaks labeled by TF name, NeuN-/Olig2-TF peaks labeled by TF name, Olig2+ TF peaks labeled by TF name, and phyloP sequence conservation.

**Extended Data Fig. 8.**
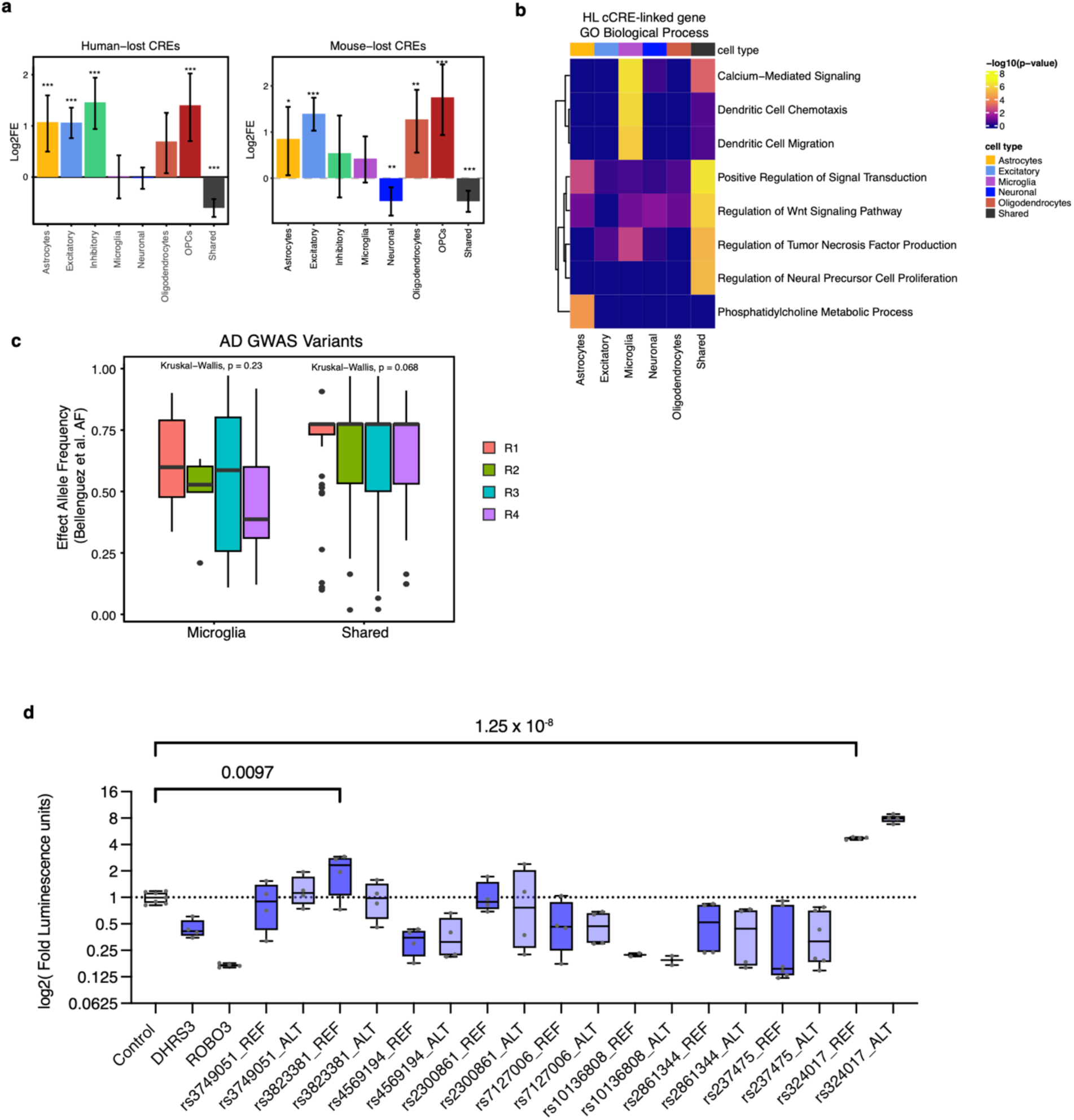
**a)** Enrichment for HL (left) and mouse-lost (right) cCREs in each cell type. Asterisks indicate Bonferroni adjusted p-values from Chi-squared test (*p < 0.05, **p < 0.01, ***p < 0.001). **b)** GO biological process enrichment using enrichR for genes linked to HL cCREs within cell type-specificity categories. **c**) AF of AD GWAS variants using frequencies observed in Bellenguez et al. **d**) Luciferase assay results for all tested variants showing the impact of schizophrenia, bipolar disorder, and neuroticism GWAS variants on enhancer activity. Labeled p-values for comparisons with a significant coefficient in the linear model.

